# Motor abstraction training generalizes to the refinement of specific movement patterns

**DOI:** 10.64898/2026.05.05.722946

**Authors:** Zekun Sun, Zhiran Xie, Samuel D. McDougle

## Abstract

The ability to abstract the invariant structure of an experience while discarding incidental details underlies generalization across virtually every domain of human cognition, from vision and language to concept learning. Yet whether the motor system generates such abstractions and whether they causally contribute to skill learning remain open questions. Here, we introduce a paradigm in which human participants learned to refine novel movement patterns by learning to precisely copy unfamiliar handwritten characters. To examine the role of motor abstractions in this form of motor learning, participants were trained on markedly rotated versions of the characters, which recruited substantially different lower-level movements while still maintaining the relevant abstract movement trajectory. Across eleven experiments, abstraction training drove robust skill improvements that were comparable to having repetitive practice on the canonical form of each character. Moreover, this learning was largely motoric in nature: it did not require a visual template, sustained trajectory feedback, or mental imagery, and was sensitive to the sequential structure of the movement trajectory. These findings establish a causal role for abstractions in motor learning, revealing that the motor system likely deploys abstractions in the earliest stages of skill acquisition.

---

Abstraction is a hallmark of intelligent systems. In vision, the brain encodes objects independent of viewpoint, lighting, and scale. In language, the same meaning can be expressed across an infinite variety of surface forms. And in concept learning, the mind extracts invariant structure from variable exemplars, enabling generalization beyond what has been directly experienced. Across these domains, abstraction — the capacity to extract the structure of an experience while discarding its incidental particulars — appears to be a core organizing principle of cognition^1–4^.

But does the motor system, which is typically cast as a non-cognitive output system, also deploy abstractions? Classic demonstrations of “motor equivalence” — the ability to produce the same action across radically different effectors and contexts, such as writing one’s signature with the dominant hand, the non-dominant hand, or even a pen held in the mouth — suggest that the motor system stores high-level features of actions that go beyond the low-level particulars of muscle commands^5–8^. Neuroimaging studies have further identified effector-independent and scale-invariant representations (e.g., writing a small letter “a” on a piece of paper versus a large one on a whiteboard) in sensorimotor cortex^5,7,9–12^. More recent studies have identified neural signatures in the sensorimotor cortex that appear to encode stroke primitives (i.e., spatio-temporal trajectories, such as a straight line or a “U” shape, but invariant to low-level features) in both nonhuman primates^13^ and humans^14^. Collectively, these findings raise the provocative possibility that the motor system itself stores, and uses, abstract features of movements.

However, the existing evidence leaves two fundamental questions unresolved. First, the abstractions implicated in motor equivalence may not be genuinely motoric in origin; it is possible that the motor system simply carries out abstraction-driven instructions inherited from other cognitive domains, translating visual, spatial, or conceptual representations into action. For example, writing one’s signature with the non-dominant hand could reflect the deployment of an over-learned visual or spatial template. The traditional theory of the generalized motor program (GMP) proposes that a class of actions is stored in long-term memory as a “program” (analogous to a computer program) that specifies the invariant features of the relevant movement sequence. When a motor program is executed, its parameters, such as amplitude, speed, or even the effector used, can be flexibly chosen to fit the context^15–18^. Such theories, however, do not make specific commitments to the nature of motor abstractions; are they motoric, mnemonic or perceptual/cognitive representations?

Second, and more importantly, classic demonstrations of motor equivalence involve overlearned, highly practiced actions^19–27^. Thus, it is not clear if behavioral reflections of motor abstraction are simply a product of long-term training or whether they play a causal role in learning, perhaps by scaffolding the acquisition of new motor skills. Answering these two questions, about the format of motor abstractions and their relevance to learning, can tell us if abstraction might be a more fundamental computational principle of the motor system. Here, we introduce a visuomotor task in which participants learned to copy novel motor patterns (i.e., unfamiliar handwritten characters) under conditions designed to isolate potential contributions of motor abstraction to movement refinement. Rather than practicing the canonical form of each movement pattern directly, we trained participants on versions of the characters rotated by extreme angles (i.e., 90°, 180°, or 270°), a manipulation that recruits substantially different lower-level movements — frequently opposite or uncorrelated in direction (Fig. 2**a** and S1) — and dramatically alters both the spatial layout of the movements and the observed visual feedback, while maintaining abstract structure. Critically, this training cannot effectively improve performance through the refinement of specific muscle commands; the primary path to improvement is through refining an abstract representation of the movement trajectory that generalizes across orientations. This rotation manipulation is a qualitatively more demanding transformation than those previously studied (e.g., scale): Rotation alters the direction of every constituent movement segment and the global spatial organization of the movement trajectory, a disruption that, in the perceptual domain, poses a well-known challenge for computational models of generalizable visual recognition^28–30^. Thus, in the logic of our paradigm, generalization of learning from a rotated to canonical form constitutes evidence that motor abstractions can contribute causally to movement refinement. Here, we also systematically tested whether the benefits of abstraction training could be explained by visual exposure, generic motor practice, mental imagery, or persistent visual feedback, ruling out each alternative in turn.

We report three main findings. First, abstraction training drove reliable improvements in the refinement of novel motor patterns, gains that were comparable in magnitude to direct repetitive practice of the canonical movement forms (Experiments 1–3). Second, the format of the abstractions being trained appeared to be primarily motoric: The abstraction training benefit was independent of mental imagery (Experiment 4), visual experience (Experiment 5), or generic motor practice (Experiment 6), and training that preserved movement trajectories but removed the visual template and sustained trajectory feedback was not only effective, but produced the strongest learning effects we observed (Experiment 7). Third, motor abstractions were sensitive to sequential structure: Reversing the temporal order of abstracted training movements significantly attenuated learning (Experiment 8). Finally, the trained abstraction proved effector independent: its benefit transferred even when training and test movements were produced by different hands (Experiments 9–11). Collectively, these findings establish that motor abstraction is neither merely a byproduct of extensive practice nor a projection of non-motor cognitive processes, but rather an intrinsic feature of the motor system that operates from the earliest stages of skill learning.

## Results

### Movement pattern refinement via motor abstraction training

If motor abstractions causally contribute to refining lower-level movement patterns, training the abstraction itself — but not lower-level muscle commands or visual templates — should lead to improved performance. To test this, we compiled 45 novel motor patterns (from the Omniglot database) for use in a naturalistic, novel handwriting task. We selected motor patterns that required a single, continuous movement (Fig. 2**a**). In Experiment 1, participants (*n* = 30) attempted to precisely copy 30 randomly-selected motor patterns by completing 30 five-trial practice blocks. In each practice block, participants first viewed and then attempted to copy a target motor pattern in its canonical (upright) orientation. During viewing, participants observed a computerized drawing of the target motor pattern from start to finish, with the viewing demos lasting on average 4 seconds, after which they were asked to reproduce the drawing to the best of their ability (Fig. 1**c**). The target motor patterns were always present on the screen during viewing and copying and thus incurred no memory demand (see Methods for further details). Next, participants completed three abstraction training trials, where the motor pattern was rotated by one of three extreme angles (90°, 180°, 270°; Fig. 1**b**) on each trial and participants attempted to copy it as before (See Methods and Fig. S1 for kinematic simulations demonstrating how the rotation manipulation alters required movement commands). After this brief training, participants again attempted to copy the motor pattern in its canonical orientation, allowing us to ask if rotation training would lead to refinement of the upright, canonical motor patterns, even though rotation training required significantly different lower-level muscle commands. From the participant’s perspective, they were simply given a long sequence of motor patterns with varied orientations to copy as best they could, without any task demands or explicit instructions to make specific improvements or attend to particular features.

**Figure 1:**
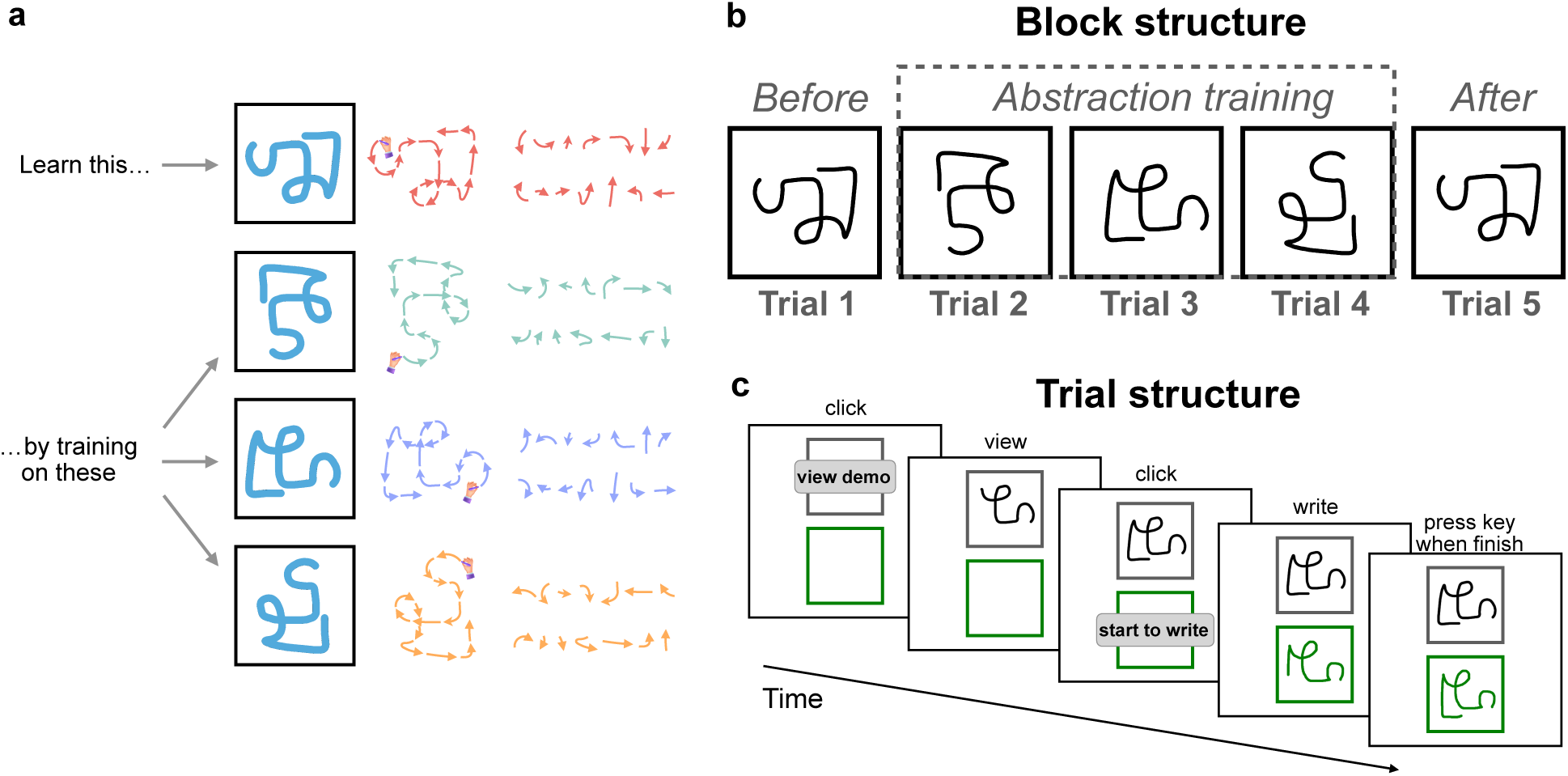
A handwriting paradigm was designed to test the role of motor abstractions in movement refinement. (**a**) Illustration of abstraction training. Participants learned to write novel patterns by being trained on their rotated versions. Rotating a motor pattern by large angles dramatically changes each step of the movement sequence.(**b**) The structure of a single five-trial practice block. In each block, participants first attempted to copy the target motor pattern in its canonical, upright orientation (Trial 1; pre-training test), then completed three consecutive training trials under the assigned training condition (Trials 2–4), and finally copied the canonical pattern a second time (Trial 5; post-training test). The primary measure was the change in performance from Trial 1 to Trial 5. (**c**) The procedure within a single trial. At the beginning of each trial, participants viewed a computerised animation of the target motor pattern being drawn from start to finish, after which they reproduced the pattern as accurately as possible with a trackpad or mouse. The target pattern remained visible on the screen throughout both the viewing and copying phases, imposing no memory demands on the participant.

**Figure 2:**
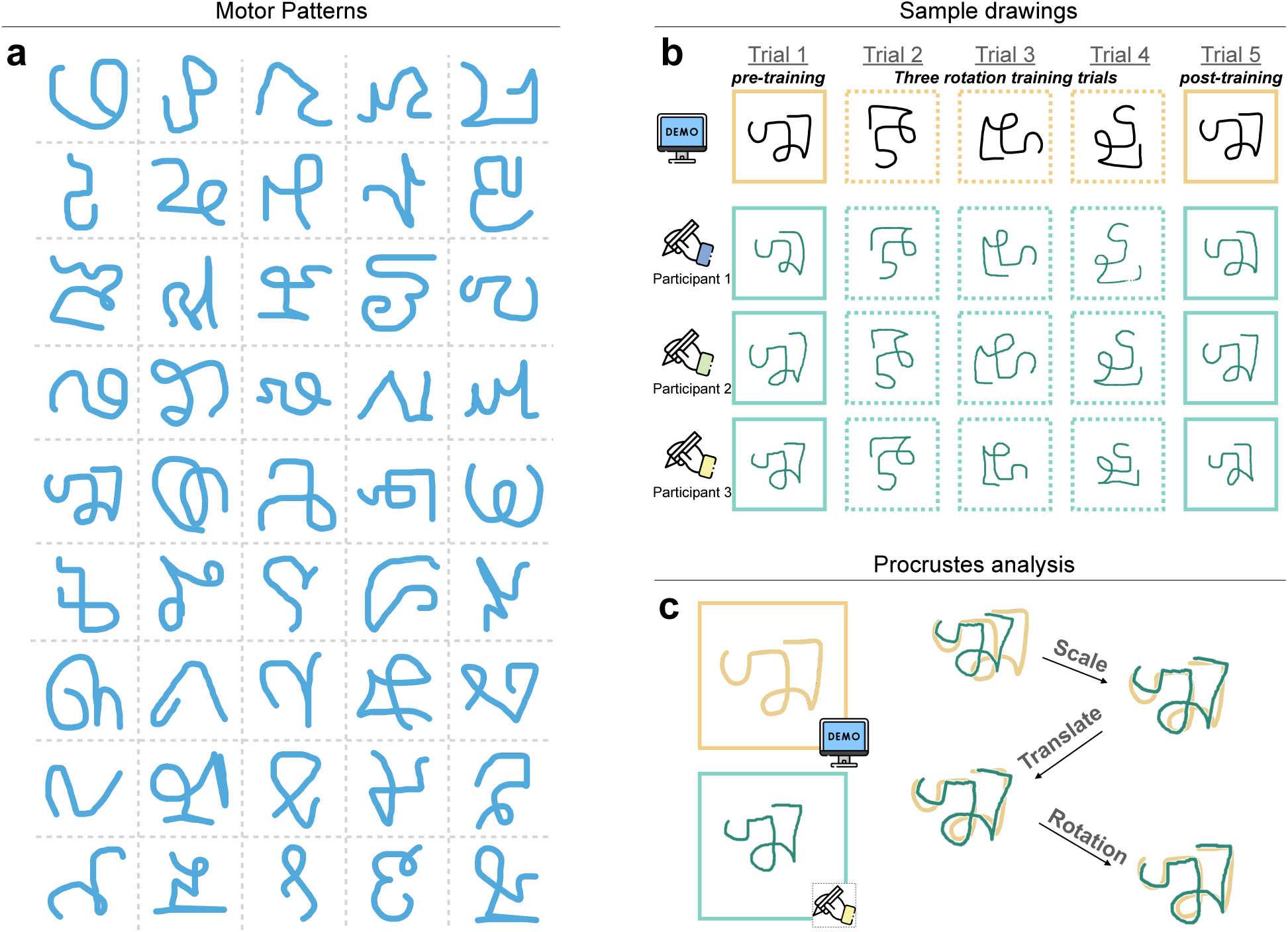
Procrustes analysis captures trajectory-level precision in novel motor pattern copying. (**a**) The 45 novel cursive motor patterns, drawn from the Omniglot database, that participants learned to copy across all experiments. Each pattern was constrained to involve a single, continuous drawing movement, ensuring a well-defined trajectory structure amenable to quantitative analysis. Participants in each experiment were assigned a random subset of these patterns (30 patterns per participant in Experiments 1–3; 10 patterns in Experiment 4–11). (**b**)Example drawings produced by three participants completing the same single practice block with one target motor pattern. The example drawings illustrate the range of initial performance across individuals from Trial 1 to Trial 5 within the block. (**c**) The Procrustes analysis procedure used to quantify copying accuracy. The algorithm aligns a participant’s drawn trajectory with the target motor pattern by optimizing for position, scale, and rotation transformations, then computes a residual shape-dissimilarity score — the Procrustes distance (*D*_proc_) — reflecting how closely the drawn trajectory matched the target. Lower *D*_proc_ values indicate more accurate reproduction. A combined performance index (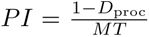, where *MT* is movement time in seconds) integrated both trajectory accuracy and writing speed, and served as the primary performance measure across all experiments. Improvements in PI therefore capture the classic dual signature of motor skill acquisition: simultaneous gains in speed and precision.

Fig. 2**b** depicts example behavior from three participants performing a single practice block with one of the motor patterns. Participants’ handwriting performance was quantified by computing both copying speed (i.e., total duration of writing) and error (i.e., the Procrustes distance [*D*_proc_], a measure of the overall trajectory dissimilarity between the target motor pattern and the copy attempt (lower values indicate more accurate copying^12,31,32^; Fig. 2**c**; see Methods). We used speed and precision because improvements in both dimensions constitute the classic signature of skill learning. Moreover, we computed a combined “performance index” measure, which was additionally used to quantify performance by taking the ratio of precision and movement time:

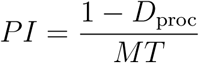

We compared participants’ performance between the first and the last trial of the practice blocks (i.e., the two upright trials). Participants’ performance copying the motor patterns at their canonical orientations significantly improved following abstraction training (*t*(29) = 5.06, *p* = 2.12 × 10*^−^*^5^, *d* = 0.92, *M_diff_* = 0.0087, 95%CI [0.0053, 0.0121]). Because each participant contributed many observations across blocks, we also fit the data with a linear mixed-effects model, including random intercepts for participant and motor pattern and fixed effects of trial (pre vs. post) and block number. This analysis confirmed a significant pre-to-post improvement following abstraction training while accounting for participant-and pattern-level variability and controlling for overall practice-related (or fatigue-related) changes across blocks (*b* = 0.0087, 95% CI [0.0047, 0.0127], *t* = 4.28, *p* = 1.98 × 10*^−^*^5^). Performance improvements involved increases in both speed and trajectory precision (Fig. 3**a**). Out of the 45 motor patterns, 39 motor patterns (87%) showed a positive training effect; that is, the performance index was higher in the fifth attempt versus the first attempt (see Fig. S2–3 for performance improvement across motor patterns and trials, and Fig. S4 for a visualization of improvements in specific sub-segments of each pattern).

**Figure 3:**
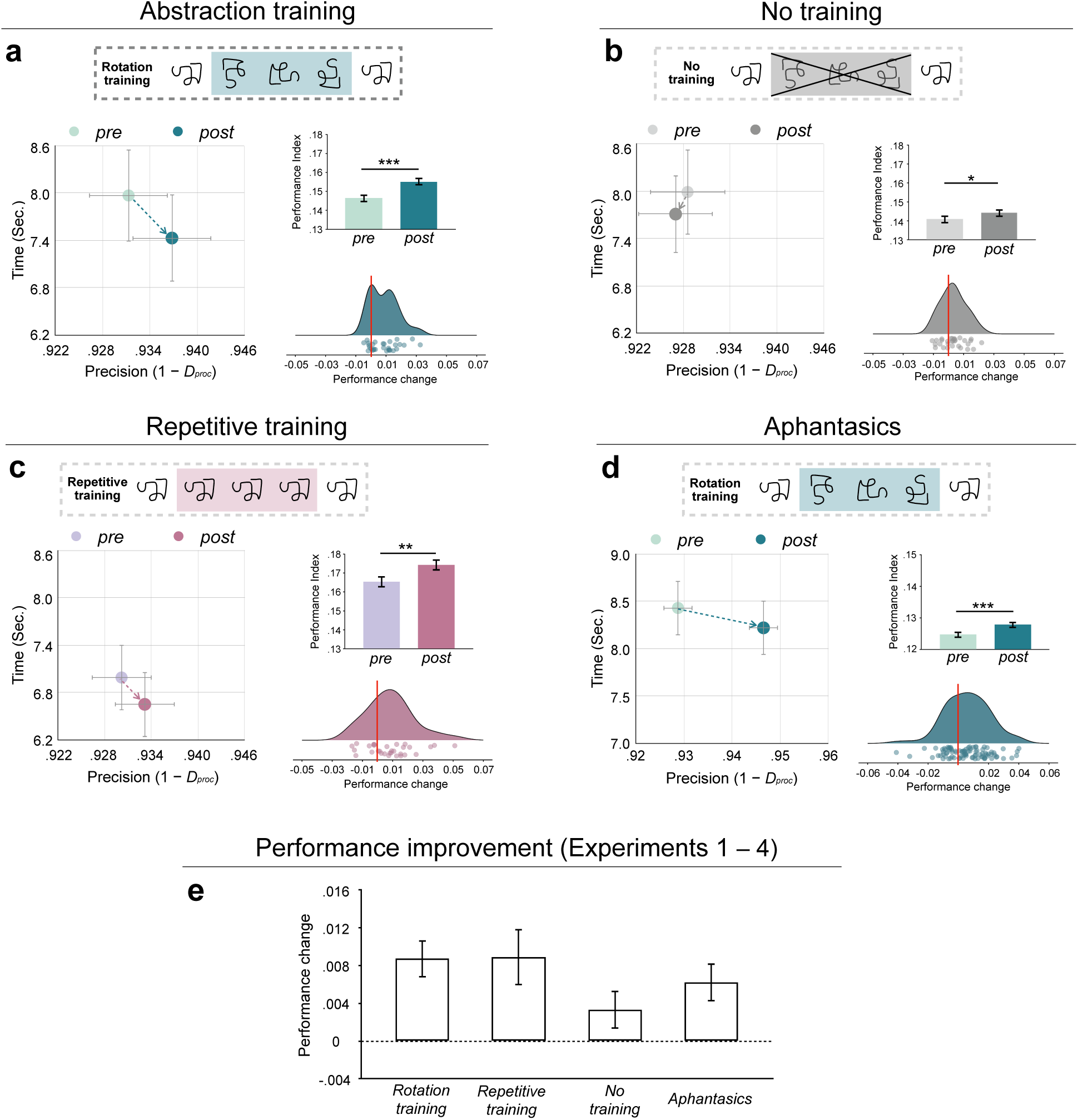
Motor abstraction training generalizes to the refinement of novel movement patterns. (**a**) The performance index (PI) on the pre-training (Trial 1) and post-training (Trial 5) test trials for participants in Experiment 1 (abstraction training; *n* = 30). During the three intervening training trials, participants copied the target motor pattern, which was rotated by 90°, 180°, or 270°. Performance improved significantly following abstraction training (*t*(29) = 5.06, *p* = 2.12 × 10^−5^, *d* = 0.92), with gains in both trajectory precision and writing speed — the dual hallmarks of skill improvement. (**b**) Pre- and post-training PIs for participants in Experiment 2 (the copy-twice control; *n* = 30), who received no training trials and simply copied the canonical motor pattern twice. Performance improved subtly (*t*(29) = 2.18, *p* = 0.037, *d* = 0.40), driven by modest increases in writing speed without corresponding gains in trajectory precision. (**c**) Pre- and post-training PIs for participants in Experiment 3 (repetitive standard copying; *n* = 30), which replaced the three rotation training trials with three additional copying attempts of the canonical, upright motor pattern. Performance improved significantly following repetitive copying (*t*(29) = 3.10, *p* = 0.0043, *d* = 0.57), with gains in both speed and trajectory precision that mirrored the pattern observed in Experiment 1. (**d**) Pre- and post-training PI for participants with self-reported aphantasia (Experiment 4; *n* = 88). Despite difficulties with conscious visual imagery, aphantasic participants showed significant improvement following abstraction training (*t*(87) = 3.93, *p* = 1.70 × 10^−4^, *d* = 0.42). (**e**) Magnitude of performance improvement (change in PI from Trial 1 to Trial 5) across Experiments 1–4. PI, performance index; *D*_proc_, Procrustes distance; *MT* , movement time; VVIQ, Vividness of Visual Imagery Questionnaire. For each experiment (a–d), the left panel plots pre- and post-training performance in the precision (1 *− D*_proc_) by movement-time space; the bar plot shows the performance index (PI) before and after training; the raincloud plot shows the distribution of per-participant performance change, with individual data points overlaid. All statistical tests in (a–d) are paired, two-sided t-tests. ^∗^*p <* 0.05, ^∗∗^*p <* 0.01, ^∗∗∗^*p <* 0.001,

We additionally analyzed the temporal coherence (i.e., the relative timing of traversing the constituent segment of each pattern) when participants copied rotated motor patterns (See details in Fig. S5); though participants’ temporal dynamics were more coherent among the copies of the rotated patterns (*r* = 0.44) compared to those of the canonical patterns (*r* = 0.13), the strength of temporal coherence did not predict the training effect (*r* = −0.017). Unlike previous work on motor equivalence^15,24,26^, this result suggests that relative timing was not be the main driving factor in abstraction training in our task (See Experiments 7 and 8 for further investigations of the putative “format” of motor abstraction).

Of course, participants could have improved simply because they wrote the canonical motor pattern twice. In a control experiment (Experiment 2) a new group of participants (*n* = 30) copied the same set of motor patterns, but they were not trained with any abstraction trials. Thus, for each motor pattern, they simply copied it twice in its canonical orientation. As expected, performance subtly improved between trials (*t*(29) = 2.18, *p* = 0.037, *d* = 0.40, *M_diff_* = 0.0033, 95%CI [0.0003, 0.0063]), with participants increasing their speed, but not precision (Fig. 3**b**). Comparing the results of Experiments 1 and 2, we found that writing improvement was over twice as large following abstraction training than after writing the motor pattern twice (0.0087 versus 0.0033, on average). A mixed-effects model (random intercepts for participant and motor pattern; fixed effects of trial and block) found that the pre-to-post improvement was not statistically reliable (*b* = 0.0033, 95% CI [*−*0.0003, 0.0069], *t* = 1.79, *p* = 0.074). Thus, the gain from copying the pattern was subtle and not fully reliable, in contrast to the robust improvements produced by abstraction training in Experiment 1 (Fig. 3**e**).

### Experiment 3: Repetitive training

The results of Experiments 1 and 2 suggest that abstraction training — writing rotated versions of a motor pattern — led to rapid movement refinement. But how does the effect of abstraction training compare to standard practice on the specific movements required for the movement patterns? To test this, a new group of participants (*n* = 30) completed three repetitive copying trials of the canonical motor patterns in place of the rotation training trials during each practice block.

As expected, participants’ performance improved significantly following repetitive standard copying (*t*(29) = 3.10, *p* = 0.0043, *d* = 0.57, *M_diff_* = 0.0089, 95%CI [0.0029, 0.0149]), with gains in both writing speed and trajectory precision. This improvement was confirmed in a mixed-effects model analysis (*b* = 0.0088, 95% CI [0.0047, 0.0130], *t* = 4.18, *p* = 3.06 × 10*^−^*^5^). The improvement following abstraction training was nearly identical in magnitude to that produced by repetitive standard copying (Rotation training versus Repetitive training; Fig. 3**e**). We quantified the relative evidence for the null using a Bayesian independent-samples *t*-test. This analysis yielded moderate evidence in favor of no difference between the two conditions (BF_01_ = 3.81), indicating that the data were roughly four times more likely under the hypothesis of equivalent improvement than under the hypothesis of a difference. This supports the conclusion that motor abstractions are sufficient to drive skill improvement in our task comparable to that produced by direct practice.

### Experiment 4: Replication with Aphantasics

Experiments 1–3 provided evidence for the effectiveness of abstraction training in movement refinement. One possible strategy in this task is to mentally simulate the movement sequence, such that the trained movements can, for instance, be mentally rotated to the canonical orientation^33,34^. Does the training effect we observed rely on some kind of conscious mental imagery? In Experiment 4, a group of aphantasics (i.e., people with a self-reported inability to visualize mental images; *n* = 88) completed a short version of Experiment 1, in which they learned to write 10 motor patterns, while also filling out the Vividness of Visual

Imagery Questionnaire (VVIQ)^35^. Though these participants reported markedly high VVIQ scores (mean score = 77, SD = 7.8, on the standard 16 – 80 total scale; higher scores indicate weaker imagery), their performance benefited from abstraction training trials (*t*(87) = 3.93, *p* = 1.70 × 10*^−^*^4^, *d* = 0.42, *M_diff_* = 0.0062, 95%CI [0.0030, 0.0094]; Fig. 3**d**; see Fig S6 for the performance across trials). The effect likewise held in a mixed-effects model analysis (*b* = 0.0061, 95% CI [0.0028, 0.0093], *t* = 3.68, *p* = 2.43 × 10*^−^*^4^). This result suggests that, at least among individuals with severe deficits in voluntary visual imagery, the benefits of motor abstraction training do not appear to require explicit mental imagery.

### Experiment 5: Visual control

People often try to learn a novel skill by first observing how others perform the task. For example, skilled individuals can transform visual templates into actions (e.g., a talented dancer may precisely imitate novel body movements immediately after seeing them for the first time). This raises a question about our results: Does rapid improvement via abstraction training arise through a form of perceptual learning, driven by the repeated visual exposure to the motor patterns during training trials? That is, participants may have extracted visual features of the motor patterns and built increasingly precise visual templates during training, leading to improved performance across trials. To test this, we ran a within-subject study in which participants (*n* = 90) either learned to draw new motor patterns via abstraction training (to replicate Experiment 1), or simply observed a computer tracing out a perfect canonical copy of the motor pattern for three trials in a row (vision training, with an attention check; see Methods). Participants completed 10 practice blocks, 5 for motor abstraction training and the other 5 for vision training (Fig. 4**a**). If the performance improvement was driven by visual training alone, performance in the vision training condition should be comparable to motor abstraction training. This was not the case: Visual blocks failed to yield a robust improvement, while we closely replicated the motor abstraction training effects in the rotation blocks (motor: *t*(89) = 4.23, *p* = 5.64 × 10*^−^*^5^, *d* = 0.45, *M_diff_* = 0.0092, 95%CI [0.0048, 0.0136]; visual: *t*(89) = 1.72, *p* = 0.088, *d* = 0.18, *M_diff_* = 0.0037, 95%CI [*−*0.0003, 0.0077]; Fig. 4**a**); the effect was significantly weakened when participants only viewed writing demos versus receiving motor abstraction training (*t*(89) = 2.14, *p* = 0.035, *d* = 0.23, *M_diff_* = 0.0058, 95%CI [0.0004, 0.0112]; Fig. 4**a**). This condition difference was corroborated by a mixed-effects model of block-wise improvements, with training condition (rotation versus vision) and block number as fixed effects and random intercepts for participant and motor pattern: learning was significantly weaker in vision than in rotation blocks (*b* = *−*0.0062, 95% CI [*−*0.0117*, −*0.0008], *t* = *−*2.24, *p* = 0.026). Thus, the abstraction training effect cannot be explained by perceptual learning or visual familiarity, and instead appears to rely on actually executing motor commands.

**Figure 4:**
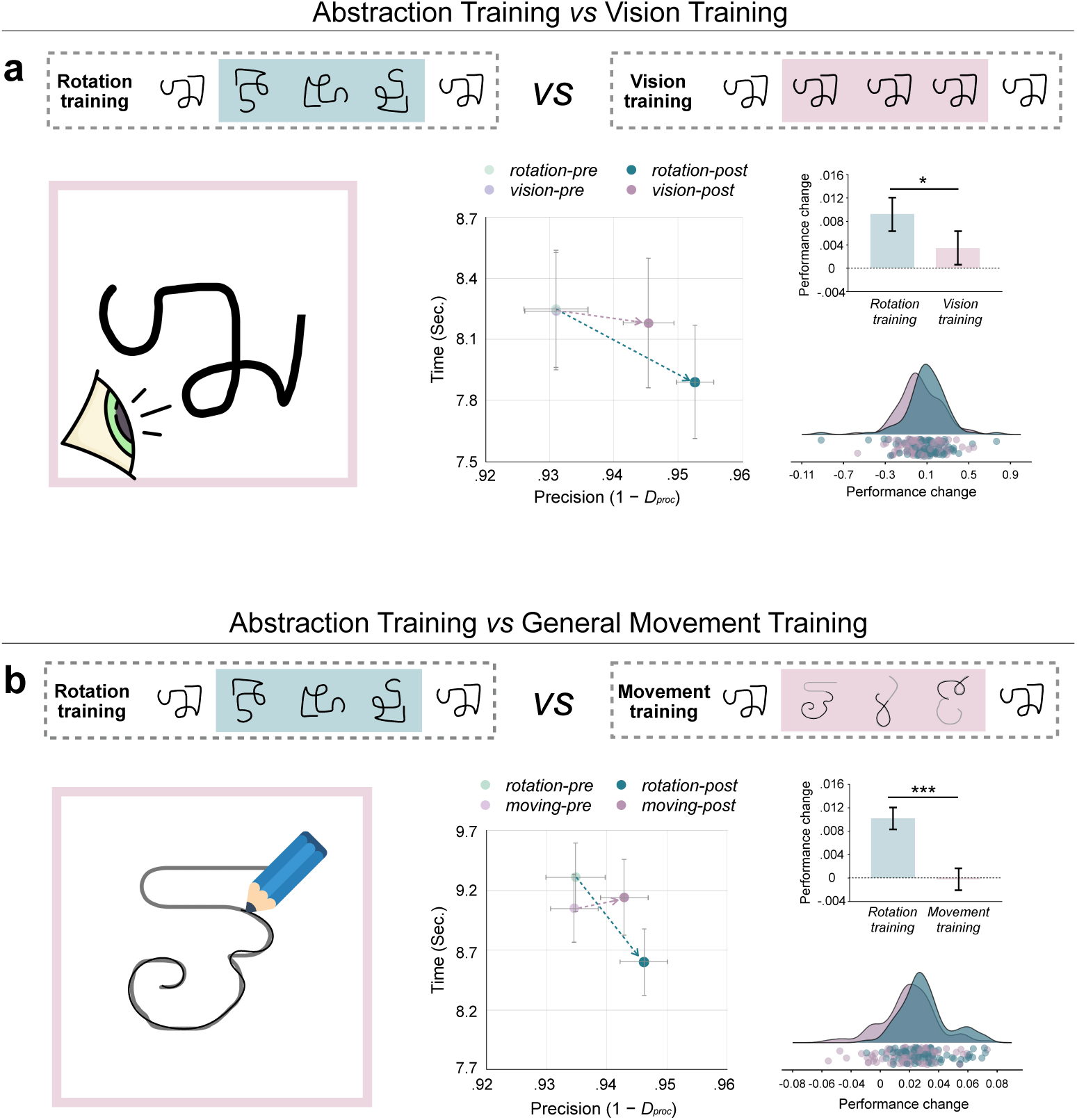
Motor abstractions cannot be explained by visual familarity or general motor practice. (**a**) Improvement across abstraction training blocks and visual training blocks in a within-subject design (Experiment 5; *n* = 90). In abstraction training blocks, participants copied the target motor pattern in rotated orientations, replicating the procedure of Experiments 1–3. In visual training blocks, participants observed a computer-generated tracing of the canonical motor pattern for three trials, with an attention check to ensure engagement. Abstraction training replicated the robust improvement observed in prior experiments (*t*(89) = 4.23, *p* = 5.64 × 10^−5^, *d* = 0.45). Visual training, by contrast, failed to produce a comparable improvement (*t*(89) = 1.72, *p* = 0.088, *d* = 0.18), and the between-condition difference was significant (*t*(89) = 2.14, *p* = 0.035, *d* = 0.23), indicating that repeated visual exposure to the target pattern is insufficient to drive movement refinement. (**b**) Performance change across abstraction training blocks and generic tracing blocks in a within-subject design (Experiment 6; *n* = 89). In generic tracing blocks, participants copied three novel motor patterns unrelated to the target pattern during the three training trials, providing general visuomotor practice but no pattern-specific motor experience. Abstraction training again produced a robust improvement (*t*(88) = 5.84, *p* = 8.77 × 10^−8^, *d* = 0.62). Generic tracing produced no detectable improvement (*t*(88) = 0.14, *p* = 0.89), and the between-condition difference was robust (*t*(88) = 4.29, *p* = 4.63 × 10^−5^, *d* = 0.45). For each experiment, the middle panel plots pre- and post-training performance in the precision (1 *− D*_proc_) by movement-time space; the bar plot shows the performance index (PI) before and after training; and the raincloud plot shows the distribution of per-participant performance change, with individual data points overlaid. All statistical tests are paired, two-sided t-tests. Error bars represent s.e.m. ^∗^*p <* 0.05, ^∗∗^*p <* 0.01, ^∗∗∗^*p <* 0.001, n.s. not significant. PI, performance index.

### Experiment 6: General movement practice control experiment

Having established that motor execution is necessary for performance improvements observed in our task, we next asked what kind of motor practice is sufficient. We first asked if people’s performance improvements were due to “generic” movement practice in the training phase (e.g., a “warming up” effect, or general short-term improvements in coordinated visuomotor actions within the task context). In Experiment 6, participants (*n* = 89) completed 5 abstraction training blocks as in the previous experiments, and in the other 5 blocks received generic motor training. The generic training consisted of three trials where participants traced the trajectory of three new motor patterns instead of the target motor pattern (Fig. 4**b**). Again, a robust improvement was found in abstraction training blocks (*t*(88) = 5.84, *p* = 8.77 × 10*^−^*^8^, *d* = 0.62, *M_diff_* = 0.010, 95%CI [0.0066, 0.0134]), replicating our previous results. However, performance did not improve in the generic tracing blocks (*t*(88) = 0.14, *p* = 0.89, *M_diff_* = *−*0.00030, 95%CI [*−*0.0046, 0.0040]), and there was a robust difference in the performance change between motor-pattern-specific abstraction training and generic motor training (*t*(88) = 4.29, *p* = 4.63 × 10*^−^*^5^, *d* = 0.45, *M_diff_* = 0.010, 95%CI [0.0052, 0.0148]). A mixed-effects model confirmed this difference: learning was significantly weaker in generic-tracing than in rotation blocks (*b* = *−*0.0113, 95% CI [*−*0.0165*, −*0.0060], *t* = *−*4.23, *p* = 2.66 × 10*^−^*^5^). Critically, this result suggests that our motor abstraction training effects reflect improvements to unique pattern-specific motor representations.

### Experiment 7: What drives the abstraction training effect?

In the next experiment we attempted to probe the “format” of the observed motor abstraction effects. Common handwriting tasks, for example, recruit overlearned visual templates and leverage online feedback control processes. However, neither observation nor online visual feedback alone seems sufficient to support ‘de novo’ skill learning (e.g., watching someone perform a backflip does not by itself enable you to execute it), suggesting a key role for movement execution in our motor abstraction effects. Next, we tested the hypothesis that the abstractions driving performance improvements in our task do not require a sustained visual template or a holistic visual representation of the pattern, but can instead be engaged through movement execution alone. That is, the benefits of abstraction training we observe may be primarily the product of the motor system executing specific movement trajectories. To test this, in Experiment 7 participants were asked to track a moving disc in each of the three training trials (see Methods). Their task in the tracking trials was simply to “chase” the target disc by following it closely with their cursor (Fig. 5a). The disc moved at a constant velocity. Unbeknownst to the participant, the target disc moved along a trajectory that exactly traced out the target motor pattern, rotated either by 90°, 180°, or 270°. If motor abstraction training effects rely on learned visual templates and/or persistent online feedback of the trajectory, the tracking task should fail to produce transfer effects. If, however, motor abstraction training improves performance primarily through the execution of the relevant movement trajectories, tracking-based training should lead to significant refinement of the canonical motor patterns.

**Figure 5:**
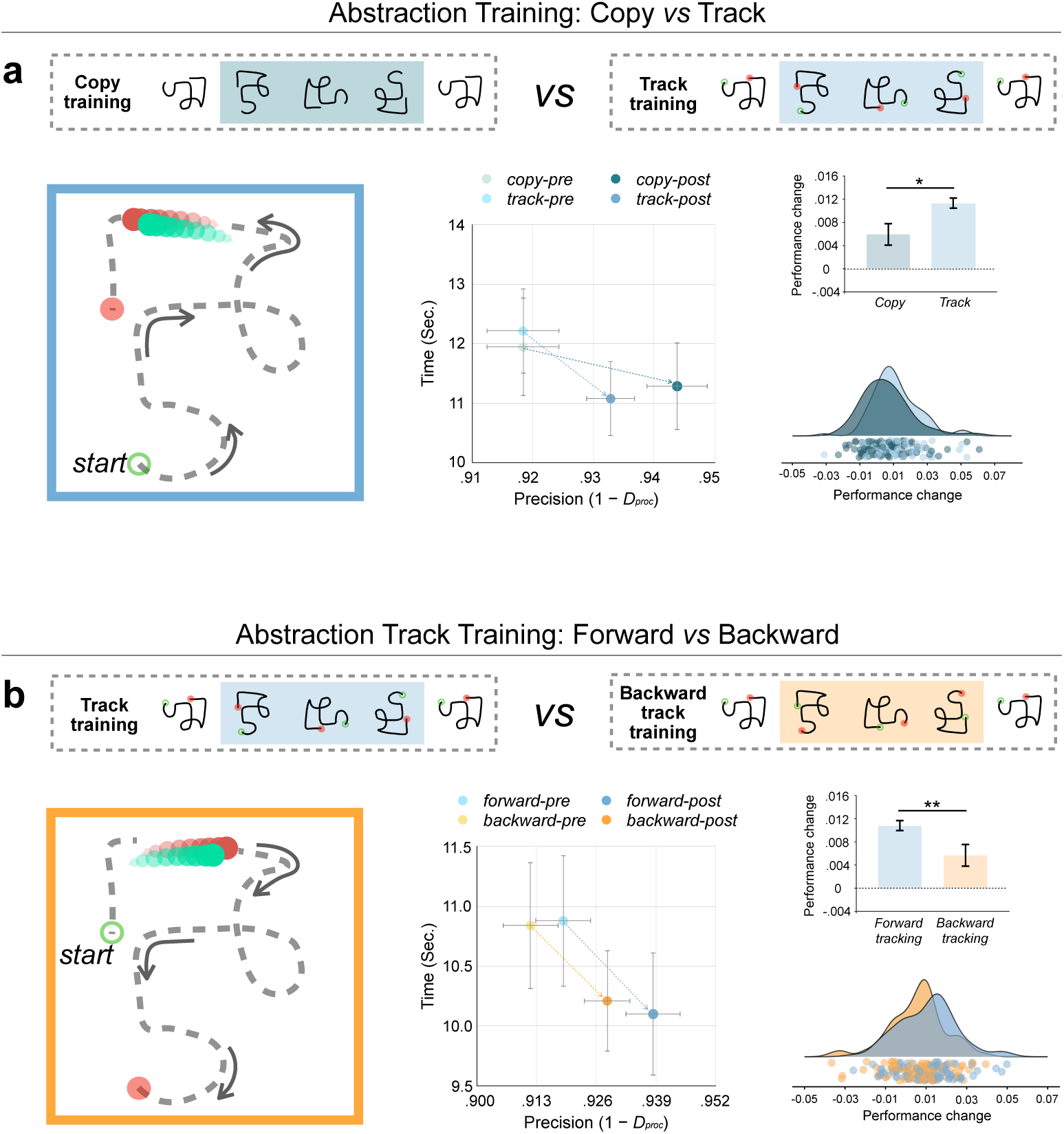
Motor abstractions are engaged through movement execution and sensitive to the sequential structure of movement trajectories. (**a**) Performance improvement across standard abstraction training blocks and tracking-based training blocks in a within-subject design (Experiment 7; *n* = 87). Standard abstraction training replicated the expected improvement (*t*(86) = 3.70, *p* = 0.00038, *d* = 0.40). Strikingly, tracking-based training produced an even stronger effect (*t*(86) = 7.68, *p* = 2.35×10^−11^, *d* = 0.82), with the between-condition difference reaching significance (*t*(86) = 2.55, *p* = 0.013, *d* = 0.27). (**b**) Forward and backward tracking training in a within-subject design (Experiment 8; *n* = 89). Both forward (*t*(88) = 7.13, *p* = 2.70 × 10^−10^, *d* = 0.76) and backward tracking-based abstraction training (*t*(88) = 3.81, *p* = 0.00025, *d* = 0.40) produced significant improvements; however, the performance improvement was approximately twice as large for forward tracking as for backward tracking (*t*(88) = 2.82, *p* = 0.0060, *d* = 0.30). For each experiment, the middle panel plots pre- and post-training performance in the precision (1 *− D*_proc_) by movement-time space; the bar plot shows the performance index (PI) before and after training; and the raincloud plot shows the distribution of per-participant performance change, with individual data points overlaid. All statistical tests are paired, two-sided t-tests. Error bars represent s.e.m. ^∗^*p <* 0.05, ^∗∗^*p <* 0.01, ^∗∗∗^*p <* 0.001, n.s. not significant. PI, performance index.

**Figure 6:**
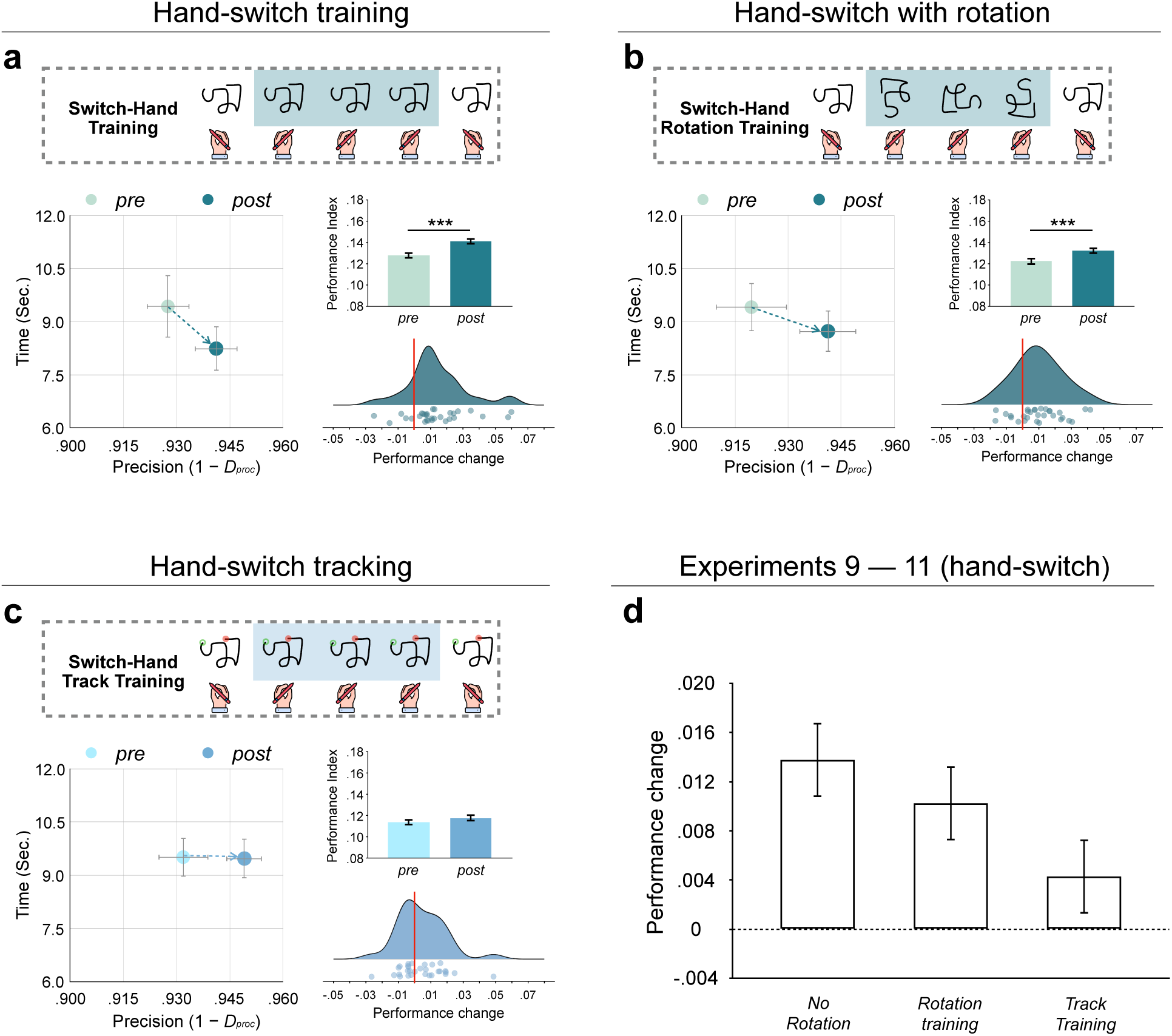
Abstraction training transfers across hands. **(a)** Hand-switch canonical training (Experiment 9), in which the three training trials were upright copies of the target pattern executed by the opposite hand. Performance on the canonical patterns improved significantly (*t*(29) = 4.11, *p* = 0.00030, *d* = 0.75). **(b)** Hand-switch rotation training (Experiment 10), in which the opposite hand copied the pattern rotated by 90^◦^, 180^◦^, or 270^◦^. Performance again improved significantly (*t*(29) = 3.72, *p* = 0.00085, *d* = 0.68), demonstrating transfer across a manipulation in which training and test share neither effector nor low-level muscle commands. **(c)** Hand-switch tracking training (Experiment 11), in which the opposite hand tracked a disc tracing the canonical pattern. Performance did not improve reliably (*t*(29) = 1.62, *p* = 0.12, *d* = 0.30). **(d)** Magnitude of the performance improvement (change in PI from pre- to post-training) across Experiments 9–11. For each experiment (a–c), the left panel plots pre- and post-training performance in the precision (1 *− D*_proc_) by movement-time space; the bar plot shows the performance index (PI) before and after training; and the raincloud plot shows the distribution of per-participant performance change, with individual data points overlaid. All statistical tests are paired, two-sided t-tests. Error bars represent s.e.m. ^∗^*p <* 0.05, ^∗∗^*p <* 0.01, ^∗∗∗^*p <* 0.001. PI, performance index.

Participants (*n* = 87) completed 5 blocks with rotation copying training (as in the previous experiments) and 5 blocks with tracking training. Performance on the canonically oriented target motor patterns improved in the standard abstraction training blocks (*t*(86) = 3.70, *p* = 0.00038, *d* = 0.40, *M_diff_* = 0.0059, 95%CI [0.0027, 0.0091]), consistent with our previous experiments. Strikingly, tracking also significantly improved performance of the canonical motor patterns (*t*(86) = 7.68, *p* = 2.35 × 10*^−^*^1^^1^, *d* = 0.82, *M_diff_* = 0.011, 95%CI [0.0080, 0.0140]), and, surprisingly, tracking-based abstraction training led to even stronger learning effects than the standard abstraction training condition (*t*(86) = 2.55, *p* = 0.013, *d* = 0.27, *M_diff_* = *−*0.0054, 95%CI [*−*0.0097*, −*0.0011]; see Fig. 5**a**, and Fig S7 for the sample tracking trajectories and the performance across trials). This advantage was reproduced in a mixed-effects model analysis, in which learning was significantly greater in tracking than in copying blocks (*b* = 0.0049, 95% CI [0.0005, 0.0094], *t* = 2.17, *p* = 0.030). Thus, we found that abstraction training without sustained visual trajectory feedback was effective at improving performance on novel motor patterns, suggesting that motor abstractions can be engaged through movement execution, even in the absence of a salient visual template, persistent trajectory feedback, or a holistic representation of the pattern.

### Experiment 8: Sequentiality in motor abstractions

Thus far, we have demonstrated that training a motor abstraction, but not lower-level muscle commands or visual templates, generalizes to improvements in performing novel movement patterns. In the next experiment (Experiment 8), we asked whether the motor abstraction constitutes a static global structure (i.e., a holistic form of the movement trajectory for each motor pattern), or if it has a temporal dimension that might reflect the sequence of sub-movements within the full movement pattern. Similar to Experiment 7, participants (*n* = 89) completed 5 tracking-based abstraction training blocks as above; however, in the other 5 blocks, they still tracked the target disc but now traced out the motor patterns from the endpoint to the start of the correct trajectory. If abstract motor representations in our task encode sequential information, then this reversed tracking condition should disrupt (or abolish) the performance improvement.

As seen in Fig. 5**b**, we observed significant skill improvement after both forward tracking and backward tracking trials (forward: *t*(88) = 7.13, *p* = 2.70 × 10^−10^, *d* = 0.76, *M_diff_* = 0.011, 95%CI [0.0081, 0.0139]; backward: *t*(88) = 3.81, *p* = 0.00025, *d* = 0.40, *M_diff_* = 0.0056, 95%CI [0.0027, 0.0085]; see also Fig. S8 for performance change for each individual motor pattern in both types of tracking training). Critically, however, this training effect was nearly twice as strong in forward tracking versus backward tracking (*t*(88) = 2.82, *p* = 0.0060, *d* = 0.30, *M_diff_* = 0.0051, 95%CI [0.0015, 0.0087]). The mixed-effects model likewise showed significantly greater learning for forward versus backward tracking (*b* = 0.0054, 95% CI [0.0008, 0.0100], *t* = 2.28, *p* = 0.023). The magnitude of the backward-tracking effect (*M*_diff_ = 0.0056) was numerically closer to that observed when participants simply copied the canonical pattern twice with no training trials at all (Experiment 2; *M*_diff_ = 0.0033). This raises the possibility that the residual improvement following backward tracking derives largely from the two canonical test trials themselves, rather than from the three intervening reverse-tracking trials. This result suggests that sequential information plays an important role in motor abstractions, further constraining the representational format of these abstractions.

### Experiments 9–11: Hand-switch generalization

Across Experiments 1–8, abstraction training transferred to the canonical form despite large changes in the required muscle commands, spatial layout, and visual feedback. In every case, however, training and test movements were produced by the same hand, which may preserve low-level command features in some cases (Fig. S1). We thus next tested effector independence, the classic hallmark of motor equivalence. Here, we report three experiments that systematically vary the content of across-hand abstraction training.

In each of Experiments 9–11, a new group of participants (*n* = 30 each) completed a short version of the paradigm with 10 motor patterns, following the block structure of Experiment 1. Within each block, participants first copied the canonical motor pattern with one hand (pre-training test), then completed three training trials with the *opposite* hand, and finally copied the canonical pattern again with the original hand (post-training test; see Methods). In Experiment 9, the three training trials were canonical copies of the target pattern (no rotation); in Experiment 10, they were the same pattern rotated by 90*^◦^*, 180*^◦^*, or 270*^◦^*; and in Experiment 11, they were tracking trials tracing out the canonical pattern (Fig. 5a–c).

Cross-hand copying produced robust transfer: Performance on the canonical patterns improved significantly following hand-switch canonical orientation training (Experiment 9; *t*(29) = 4.11, *p* = 0.00030, *d* = 0.75, *M*_diff_ = 0.014, 95% CI [0.0070, 0.0210]; Fig. 5a) and following hand-switch rotation training (Experiment 10; *t*(29) = 3.72, *p* = 0.00085, *d* = 0.68, *M*_diff_ = 0.010, 95% CI [0.0045, 0.0155]; Fig. 5b), even though the trained and tested movements were executed by different limbs. Mixed-effects models confirmed a significant improvement following cross-hand training (Experiment 9: *b* = 0.013, 95% CI [0.0069, 0.020], *t* = 4.02, *p* = 6.76 × 10*^−^*^5^; Experiment 10: *b* = 0.010, 95% CI [0.0042, 0.016], *t* = 3.37, *p* = 7.99 × 10*^−^*^4^). We note that the raw magnitude of the performance improvement (*M*_diff_) produced by rotation-based training was remarkably consistent across experiments, clustering around 0.009–0.010 in Experiments 1, 3, 5, and 6. Strikingly, the first two hand-switch studies produced improvements of comparable magnitude (*M*_diff_ = 0.014 and 0.010, for Experiments 9 and 10, respectively).

The cross-hand transfer, however, did not extend to tracking to the same degree: Following hand-switch tracking training (Experiment 11), performance on the canonical patterns did not improve reliably (*t*(29) = 1.62, *p* = 0.12, *d* = 0.30, *M*_diff_ = 0.0043, 95% CI [*−*0.0011, 0.0097]; mixed linear model: *b* = 0.0037, 95% CI [*−*0.0017, 0.0091], *t* = 1.34, *p* = 0.18; Fig. 5c). The pattern across the three experiments — robust transfer for canonical and rotated copying, but not for tracking (Fig. 5d) — suggests that the abstraction engaged by tracking, might be more closely tied to the trained effector than the abstraction engaged by copying. Overall, that rotation training transferred across hands, at a magnitude on par with same-hand training, extends our key results to a regime in which training and test share neither effector nor low-level muscle commands.

## Discussion

Abstractions play a central role in most domains of cognition and perception, but the role of abstractions in motor learning and control is unclear. Here we asked if motor abstractions causally contribute to the refinement of movements during brief practice, and what the format of these motor abstractions may be. Our findings reveal that motor abstraction is likely not simply a byproduct for expert, overlearned tasks (as demonstrated in classic textbook examples of “motor equivalence” for, e.g., handwriting). Across eleven experiments, we demonstrate that the motor system likely recruits abstractions from the earliest stages of skill learning — emerging within only a handful of practice trials, long before the extended practice typically associated with the consolidation of specific lower-level movement patterns. Participants who trained on rotated versions of novel motor patterns, which induced movements with distinct muscle commands, spatial trajectories, and timing profiles relative to the target movement patterns, nonetheless showed robust improvements in the target movements. Visual learning, generic motor practice, mental imagery, and consistent relative timing could not account for these robust transfer effects. We believe that our results reflect the causal operation of motor abstractions in motor learning. That is, motor abstractions that are invariant to specific execution parameters are leveraged by the motor system during skilled performance and learning.

The learning transfer we observed constitutes a novel form of motor generalization that differs from the typical kinds of generalization previously documented in the motor learning literature. Classical motor generalization describes transfer across effectors, size, scales, or contexts in which the training and test movements share substantial overlap at the level of intrinsic muscle commands, extrinsic spatial trajectories, or explicit timing profiles^15,20–22^. Our rotation manipulation, by contrast, alters the direction of each movement segment and the global spatial organization of the movement trajectory. Rotation is also a transformation that proves notoriously difficult for computational models of pattern recognition (e.g., in computer vision) without explicit training or specialized architectural design features^28–30^. That the motor system generalizes across this difficult transformation speaks to the flexibility of the abstract representations it appears to encode.

Our results further suggest that the format of these abstractions can be engaged through movement execution, independent of a visual template, persistent trajectory-error feedback, or perception of the holistic form of the movement goal. In Experiment 7, participants who tracked a disc moving at constant velocity along the rotated trajectory of a target motor pattern, without seeing the form of the pattern itself and without the natural acceleration-deceleration profile of real handwriting, showed robust generalization to the canonical target patterns. This finding is surprising for two reasons. First, it demonstrates that the abstractions contributing to learning may operate independently of visual templates. Second, the constant-velocity constraint removed the relative timing structure of the movement, the characteristic acceleration and deceleration profile that prior motor equivalence work has identified as a key invariant feature of generalizable motor programs^19,21,22,24,36,37^. This suggests that the motor abstraction recruited in our task is neither a visual template nor a relative timing or force time-course, but an even more abstract representation of the movement structure.

A natural next question concerns what, exactly, the motor system abstracts in tasks like ours. Our results indicate that whatever underlies the transfer is invariant to absolute spatial parameters, visual form, and low-level movement commands, yet sensitive to sequential order. These effects could be described via a stored, flexible representation of the holistic movement structure, or perhaps via incremental tuning to specific invariant properties of the movement patterns in space and time. Our main contribution here is showing that the motor system can extract the abstract structure of a movement to improve performance even across large transformations of the requisite low-level commands. Characterizing the exact nature of this abstraction is an exciting open question, and future computational work can help to adjudicate between various motor abstraction models to explain our findings.

One particular aspect of our results was quite surprising, where tracking-based training, in which participants followed a moving disc, drove equivalent or even stronger transfer effects than copying a fixed visible pattern (Experiment 7). This may seem paradoxical, since copying allows participants to view the whole pattern and construct a feedforward model of how to produce it, whereas tracking is guided by feedback control to track an external sensory target. However, we note that the holistic visual model available during rotation-copying is a model of the *rotated* form, which must then be transformed back to the canonical orientation at test, a transformation that may carry its own cognitive cost or interfere with canonical pattern production. Tracking, on the other hand, provides no such competing template and instead primarily targets the movement trajectory and somatosensory feedback. Perhaps directly practicing the movement segments is more efficient in the earliest stage of learning than practicing it in relation to a visual model. To further speculate, it may be that more purely motoric practice, with minimal visual input, is inherently more transferable, much as musical improvisation, unanchored to a score, generalizes across contexts. Testing this idea will require further work and additional manipulations of sensory feedback.

Our results also raise a further question concerning whether motor abstraction functions independent of visual and proprioceptive sensory feedback. Addressing this will require dissociating the sensory, planning, and execution-based contributions to abstraction-driven improvements. A neuropsychological approach could be useful here: Testing our task on patients with peripheral sensory neuropathy, in whom proprioceptive feedback is disrupted, could help establish the role of proprioceptive input in abstraction transfer effects; conversely, testing patients with frontoparietal lesions, such as those with ideomotor apraxia, in whom planning and imitation is disrupted despite intact sensation and movement execution, could probe the role of central, praxis-based processes. Indeed, recent work in such patients has shown that despite having normal movement execution skill, they have significant deficits imitating abstract movement trajectories^38^.

The ideas we propose here may be relevant to designing novel training paradigms that exploit abstraction-mediated generalization, by using varied, non-canonical movement experiences to potentially improve the acquisition of motor skills. Helpful practice variations may be those that alter the low-level execution of a movement while minimally disturbing its abstract, sequential structure. In rehabilitation, for example, patients recovering from stroke or injury are often limited in their ability to practice a target movement directly. Speculatively, our results suggest that practicing structurally-related variants (e.g., the same movement mirrored, rescaled, reoriented, or performed with the less-affected limb) may improve recovery of the target action even though these variants recruit different lower-level movements. In sports and musical instrument learning, where conventional wisdom emphasizes high-fidelity repetition of target movements, the same principle would suggest that practicing a stroke or a musical passage across varied tempos, amplitudes, orientations, or effectors may be especially effective at building the abstract representations that underlie skilled performance^39–42^.

Indeed, the value of variable practice is well established^43^, both in traditional accounts (e.g., schema theory, which proposes that people abstract a schema — a rule or relationship linking the initial conditions, the parameters, sensory consequences, and goal-related outcome — for a whole classes of movements^15–17^) and in contemporary “constraints”-led approaches. For example, our proposal aligns closely with “differential learning” ideas, in which highly variable, non-repetitive practice has been shown to enhance skill acquisition and retention^44,45^. But whereas differential learning emphasizes stochastic variability, our results indicate that specifically structurally-consistent variation may be most effective.

Additional key questions remain for future work. First, the neural substrates of motor abstraction in skill learning are mostly unknown. Neural data have identified effector-independent and scale-invariant representations in sensorimotor cortex for familiar or well-trained actions^10–14^, but whether these same regions support the kind of invariant abstraction we document here, and whether their engagement changes as a function of practice, remains to be fully determined. Prior work on the neural bases of sensorimotor adaptation, transfer of learning, and “learning to learn” offers a natural starting point: this literature implicates cerebellar-thalamocortical circuits in the storage and transfer of adapted movements and cortico-striatal and cingulate networks in the more cognitive components of early learning^46^; it will be important to ask whether these same systems support the pattern-specific abstraction reported in this work.

Second, our paradigm examined motor abstraction over a compressed timescale within a single experimental session, across a modest number of practice trials. This was by design, as we were primarily interested in the earliest stages of refining a novel movement pattern. But how motor abstractions evolve over longer timescales, and how they interact with the consolidation and retention of movement-specific representations following extended practice, are key open questions. These questions connect naturally to influential two-process accounts of procedural learning, in which an early, fast-evolving, effector-independent process that represents movement sequences in non-motor (e.g., visuospatial) coordinates is complemented by a slower-developing process that represents them in intrinsic, effector-specific motor coordinates^47^. Abstraction-mediated generalization may be most powerful precisely at these earliest stages, with its contribution diminishing as movement-specific, effector-specific traces consolidate. Alternatively, abstract and specific representations may co-develop and mutually constrain each other throughout learning. Our findings complicate the neat mapping between effector-independence and a non-motor visuospatial code, as the abstraction effects were effector-independent and not purely visuospatial in format.

Third, the present paradigm used a discrete, well-defined motor task: Copying novel motor patterns under non-ecological conditions. Whether abstraction-mediated generalization operates similarly in more naturalistic, continuous, or high-dimensional motor skills, such as playing a musical instrument or acquiring a sport-specific technique, remains an exciting direction for future work.

Taken together, the findings here reveal that the motor system may use abstract representations of movement patterns during learning. The format of these abstractions appears to be more motor than visual, and the representations appear to be both flexible (e.g., invariant to absolute spatial or temporal parameters and effector independent) and constrained (e.g., sensitive to sequential structure). We propose that the motor system can deploy abstract structures in the service of refining concrete (muscle-specific) movements. Much as evolving traditions in the study of vision have re-framed the visual system as genuinely “high-level” in many key ways (e.g., in that it satisfies key cognitive desiderata like abstract, compositional, and invariant representation^48–52^), the present findings, together with other previous and emerging work^13,34,38,53–55^, invite two complementary readings. On one hand, they reveal a high-level, cognitive function putatively operating within the motor system, casting the motor system as surprisingly cognitive in its architecture. On the other hand, the results could be said to point in the opposite direction: if abstraction is partly a motor process, then perhaps analogous functions in cognitive domains reflect an extension of action-oriented computations. This possibility resonates with a longstanding tradition that views higher cognitive capacities as being organized primarily around action^56–59^. Either way, our findings support the idea that the boundary between “motor” and “cognitive” processes is more porous than traditionally assumed.

## Methods

### General methods

All protocols were approved by Yale University’s Institutional Review Board, protocol number 2000027351. Experiments 1–4 were not pre-registered. For Experiments 5–8, we pre-registered the sample size, experimental design, and main statistical tests. We report all of these pre-registered analyses; in addition, we also report additional follow-up analyses to more holistically model the effects (such as using mixed-effects models that include random-effects terms for participants and motor patterns). All experiments were coded using a combination of Hypertext Markup Language (HTML), Cascading Style Sheets (CSS), and JavaScript (JS). The data, experiment code, stimuli, and (where applicable) pre-registrations for all studies are available at (https://osf.io/2gw5c). Readers can also experience example trials of all of our experiments at (https://zk.actlabresearch.org/abstraction).

### Participants

We recruited all participants (except for the aphantasic sample) via the online platform Prolific (total N = 490; https://www.prolific.com/); for a discussion of the reliability of this participant pool, see^60^). For Experiments 1–3, we recruited 30 participants each. For Experiment 4, aphantasic participants were recruited via the Aphantasia Network (see below). For Experiments 5–8, we recruited 100 participants each. Each individual participant was recruited only for one experiment. Participants were pre-screened to be between 18–35 years of age, have a minimum approval rate of 99%, at least 50 prior submissions, normal or corrected-to-normal vision, fluency in English, and U.S. residence. Aphantasic participants (Experiment 4) were recruited from the Aphantasia Network (https://aphantasia.com/), through which our web experiment was distributed via an email link, such that participants could complete the task along with the VVIQ questionnaire. In total, 93 aphantasics participated in that study. All participants in these experiments provided informed consent for their participation.

### Motor patterns

We selected 45 novel motor patterns from rare languages in the Omniglot data set. The standard motor patterns for participants to imitate were actual handwriting samples from online writers collected in an earlier study^61^. We mirror-flipped the motor patterns horizontally such that the motor patterns used here would not be found in any existing written languages. We selected these writings based on several criteria: (1) they were single-stroke motor patterns; (2) they were not similar to commonly-used letters or symbols (e.g., English or Greek); and (3) they were legibly written.

In Experiments 1–3, 30 motor patterns were randomly selected from 45 motor patterns for each participant. In Experiments 4–8, participants were trained on the same 10 out of 45 motor patterns, which had induced reliable training effects in the first experiment.

### Kinematic simulations

To estimate how different the rotated training patterns were in their lower-level kinematics versus their canonical counterparts, we constructed two planar inverse-kinematic models corresponding to the two input devices used by participants (Fig. S1). The mouse model represented drawing as a shoulder–elbow–palm kinematic chain. With shoulder position fixed, it tracked shoulder and elbow angles. The trackpad model represented drawing as an elbow–wrist–fingertip kinematic chain (similar to finger painting). With elbow position fixed and finger curl constrained, it tracked forearm and wrist angles. For each of the 45 motor patterns, the template cursor trajectory was mapped to physical units, anchored at the starting point as the origin, and rotated to simulate the 90*^◦^*, 180*^◦^*, and 270*^◦^* orientations we tested. Closed-form inverse kinematics converted these cursor paths into joint-angle time courses. For each joint, we then computed the cosine similarity between the canonical time course and each rotated one.

Across patterns, drawing the rotated forms involved joint movements that were either opposite in direction or significantly uncorrelated with those of the canonical form, supporting our assumption that the rotation manipulation substantially altered the lower-level movements required (Fig. S1). Full documentation of both models, including per-pattern results, is available at https://zk.actlabresearch.org/abstraction/movement_models.html

### Experiment 1: Rotation training

#### Design and procedure

The task consisted of 30 blocks, and each block contained 5 trials, resulting in 150 writing trials in total. The standard (canonical) motor pattern was shown in the first and last trials of a block. Between these two testing trials were three training trials, in which participants copied the motor pattern rotated by three extreme angles — 90°, 180°, 270° (in this fixed order; 1**b**). At the start of each trial, participants saw two boxes vertically aligned at the center of their screen, with a “view demo” button shown in the box on the top (the demo box). Participants clicked the button to view an animation demonstrating how to write the motor pattern. Though participants may learn to write by just seeing a static image of the motor pattern, we asked participants to watch the writing demo for several reasons: (1) it instructed participants on the exact writing sequence in a continuous movement (i.e., where to start); (2) it standardized the learning process across participants, such that all participants observed how to write the target motor pattern for the same length of time before copying it themselves; and (3) it showed the detailed features and movement of the canonical pattern. Participants’ cursor was hidden once they clicked the “view demo” button. After the demo was done, a “start to write” button appeared in the box below (the writing box). When participants were ready to write, they clicked on the writing button and were given the starting point for writing while their cursor was again hidden. Next, they simply moved their mouse or finger on their trackpad to reproduce the same movement pattern as they saw in the demo to the best of their ability. Once they finished writing, they pressed the space-bar to proceed to the next writing trial (see 1**c** for an example writing trial).

At the beginning of the writing task, participants were told that all motor patterns should be completed with a single, continuous movement and that they had no chance to rewrite or edit their responses. A scoring sheet was given to the participants (Fig. S9) to instruct them to copy the exactly same motor patterns without changing or omitting any features. To motivate participants to do their best, they were told that the top writers would receive a monetary reward afterwards — we assigned a bonus to the top 10% of writers based on their writing accuracy in each experiment. Participants were not informed at the outset about the essential design choices of the task, such as the practice blocks with testing and training trials; from their perspective, the goal was to simply copy a series of motor patterns as accurately as possible.

The demo and writing boxes were displayed in participants’ web browsers at the size of 300 × 300 pixels. No time pressure was applied to participants’ writing. The order of the 30 blocks was randomized across participants.

### Analyses

On each trial, we recorded participants’ writing trajectory and movement duration. Participants’ writing performance was evaluated by their copying speed and error. Here, speed is the total time taken to complete a motor pattern. Error was operationalized as the Procrustes distance between a target motor pattern and the attempted copy. Procrustes analysis quantifies the distance or dissimilarity between two shapes (or patterns of points) by minimizing the distance between them after they have been optimally aligned through translation, rotation, and scaling. To calculate Procrustes distances, all motor patterns and their copies were resampled to 1,000 consecutive points at uniform intervals using linear interpolation. The Procrustes distance metric ranges from 0–1, where larger numbers indicate higher dissimilarity between two shapes (i.e., more error, or lower accuracy in copying the shape of the target motor pattern). Speed and error can be traded off in writing, so we additionally computed a performance index (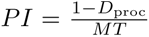) to measure changes in writing skill. The index can be understood as copying precision (1 *− D*_proc_) per unit of time (1 second). The performance change (i.e., the training effect) was the difference in PI between the pre-training trial and the post-training trial, calculated for each practice block. An increased PI value after training is observed when participants (a) drew a more accurate copy within a similar amount of time; (b) completed a similarly accurate copy in a shorter time; or (c) improved in both precision and speed.

No participants were excluded. On the trial level, we excluded any trials with missing values and/or extremely large Procrustes distances — indicating no apparent intention to copy the target motor pattern — defined as more than 3 standard deviations above the mean Procrustes distance. We also excluded a full block if either the pre-training or post-training trial was flagged as an outlier. On average, approximately 8 out of 150 trials (5%) were excluded per participant. We ran a two-tailed, paired *t*-test (*α* = .05) to compare participants’ pre-training performance with their post-training performance. No results reported here depended on excluding or keeping these trials.

As expected, participants had little difficulty copying the motor patterns, with a mean Procrustes distance of 0.075 and a mean completion time of 7.7 s. Their copies also accurately reflected the right orientation and size of target motor patterns. Thus, our task tapped into a well-learned adult skill — visuomotor copying of visible shapes. On average, writing was scaled down to 93% and rotated by 0.04° to match the target motor patterns (i.e., the optimal scaling and rotation for Procrustes superimposition of the copy to the target motor pattern). In addition, there was no significant difference in specific Procrustes transformations between pre- and post-training trials, suggesting that any improvement in writing accuracy and speed was not caused by copying a motor pattern in a different size or rotation. The overall performance across five trials and participants in the practice block is shown in Fig. S3.

To complement the pre-registered *t*-tests, and following a reviewer’s suggestion, we additionally fit each experiment’s data with a linear mixed-effects model, with degrees of freedom and *p*-values estimated via Satterthwaite’s method. For Experiment 1, performance index was modeled as a function of trial (pre vs. post) and block number, with random intercepts for subject and motor pattern.

### Experiment 2: No training control

Experiment 2 examined whether participants’ performance improved simply because they wrote the same standard motor pattern for the second time rather than because they had been trained via the rotated versions. We instructed participants to copy 30 randomly chosen motor patterns, copying each motor pattern in its canonical orientation twice in two consecutive trials, resulting in 60 trials in total. Similar to Experiment 1, we excluded trials with extremely large error (Procrustes distance *>* 3 standard deviations), which accounted for approximately 5% of trials per participant on average. On average, specific Procrustes transformations scaled down hand copies by 93% and rotated them by 0.08°. Neither Experiment 1 nor Experiment 2 showed evidence for meta-learning — the training effects did not increase in magnitude as the experiment proceeded. As in Experiment 1, we also fit a linear mixed-effects model to the corresponding data.

### Experiment 3: Repetitive copying control

Experiment 3 addressed the possibility that abstraction-driven improvements could reflect repetitive motor practice of any kind. We recruited 30 participants, independent of those in Experiments 1–2. The design was identical to Experiment 1 in all respects, with one exception: the three training trials used the canonical, non-rotated motor pattern instead of rotated versions. Participants completed the 30 blocks with the same stimuli selection procedure as Experiment 1. All other procedures and exclusion criteria were identical to Experiment 1. No participants were excluded. As in Experiments 1 and 2, we also fit a linear mixed-effects model to the corresponding data.

To compare the training effects between Experiments 1 and 3, we used a Bayesian independent-samples *t*-test to quantify the relative evidence for the null (no difference) against the alternative (a difference), for the comparison between abstraction training (Experiment 1) and repetitive standard copying (Experiment 3). Analyses were conducted with the BayesFactor package in R, using the default Jeffreys–Zellner–Siow prior — a Cauchy distribution on the standardized effect size with scale *r* = 0.707. We report BF_01_, the ratio of the marginal likelihood of the data under the null to that under the alternative, with values above 3 conventionally interpreted as moderate evidence for the null. To confirm that this conclusion did not depend on the width of the prior, we repeated the analysis with wider prior scales (*r* = 1.0 and *r* = 1.414), which yielded the same qualitative conclusion.

### Experiment 4: Aphantasics

Experiment 4 tested whether rotation training improved writing skills in participants with aphantasia — individuals who report a complete or near-complete absence of voluntary visual mental imagery. The motor patterns and procedure were similar to Experiment 1 (with some minor changes identical to those used in Experiments 5–6; see below). Participants completed 10 rotation training blocks. The web link to our task was distributed through the Aphantasia Network, a specialized community of individuals who self-identify as having little or no visual imagery. Participants voluntarily joined the experiment and completed the Vividness of Visual Imagery Questionnaire (VVIQ) during a three-month time window. In total, 93 participants completed the task. Five participants were excluded for failing to provide a complete data set, leaving 88 participants with analyzable data. We scored the VVIQ on its standard 16–80 total scale, on which higher scores indicate weaker imagery and the ceiling of the scale corresponds to a near-complete absence of voluntary visual imagery. Participants had a mean total score of 77 (SD = 7.8), with 88% scoring above 75 — near the ceiling of the scale — confirming that the great majority experienced severe imagery impairment. Because the sample was recruited specifically through the Aphantasia Network and was overwhelmingly composed of severely impaired imagers, we analyzed the group as a whole rather than restricting to a subset. As before, trials with extremely large error were excluded. As in Experiments 1-3, we also fit a linear mixed-effects model to the corresponding data.

### Experiment 5: Vision control

#### Design and procedure

This experiment examined whether the improvement in writing was driven by visual training. Ten motor patterns were selected from previous experiments and used here. Participants learned to write 5 motor patterns with rotation training, as in Experiment 1. In addition, they learned the other 5 motor patterns with vision training. In a vision block, participants first copied the standard motor pattern. Then, they had three vision training trials, in which the program reproduced a perfect copy for them after the demo video. In vision trials, participants were asked to press the spacebar immediately after the program completed the copy, as fast as possible (to insure attention was paid). Finally, they copied the motor pattern again. The motor patterns for rotation versus vision blocks were randomly assigned for each participant, and the order of blocks was randomized.

### Analysis

As stated in our pre-registration, we recruited 100 participants for this experiment, and each completed 10 writing blocks (50 trials). On the trial level, we excluded those with extremely large Procrustes distances or missing values, as described above. For vision training, a trial was excluded if participants pressed the attention check key >3 seconds after the observed writing was complete. At the participant level, those who had >15% of invalid attention check trials, reported technical problems, or did not provide complete data were excluded. Three participants were excluded for failing to provide a complete data set and 7 for having over 15% of trials with large error, leaving 90 participants with analyzable data. For each within-subject Experiment (Experiments 5–8), we fit a mixed-effects model on block-wise PI changes as a function of training condition and block number, with random intercepts for subject and motor pattern.

### Experiment 6: General motor control

Experiment 6 examined whether the improvement in writing was driven by generic effects of coordinated visuomotor practice during training. Participants completed 10 writing blocks (50 trials), 5 rotation blocks and 5 general movement blocks. In a generic movement trial, participants first saw the writing demo of a novel motor pattern and were then asked to trace the shaded trajectory of that motor pattern. Across a generic movement block, participants first copied a target motor pattern, then traced three new motor patterns, and finally copied the target motor pattern again.

We pre-registered the same sample size and exclusion criteria as in Experiment 5. Six participants were excluded for failing to provide a complete data set and 5 for having too many invalid trials, leaving 89 participants with analyzable data.

### Experiment 7: Tracking-based training

#### Design and procedure

This experiment examined whether rotation training improved writing when participants simply tracked a target moving at constant velocity along the trajectories of rotated motor patterns, such that no visual template and no temporal properties were encoded during training trials. The same 10 motor patterns used in Experiments 4–6 were also used here. Participants learned 5 motor patterns in rotation-copy blocks, and the other 5 in rotation-tracking blocks, where they first copied a target motor pattern, then completed three tracking trials, and finally copied the target motor pattern again. A tracking trial began with a “start to track” button at the center of a blank screen, with the instruction “move the blue disc inside the red one” shown below the button. Immediately after participants clicked this button, they saw a smaller blue disc inside a larger red disc, with a three-second countdown. Once the timer reached zero, it disappeared and the red disc began to move. Participants moved the blue disc (which represented their cursor) to track the red one as closely as possible. Unbeknownst to participants, across the three tracking trials the target disc (red) moved along the trajectory of the target motor pattern rotated by three angles (90°, 180°, 270°) at uniform speed (∼20% slower than the writing demo used in copying trials, to reduce the difficulty of tracking).

### Analysis

Exclusion criteria here were the same as in Experiments 5–6: trials were excluded for large error, and participants were excluded if more than 15% of trials were invalid (9 participants) or if they did not provide complete data (4 participants). As can be seen in Fig. S7, participants’ hand movements accurately followed the shape of the rotated motor patterns, while making more local errors in tracking trials given the feedback control requirements and uncertainties of tracking.

### Experiment 8: Forward/backward tracking

Experiment 8 examined whether the specific canonical sequential order of sub-movements within each movement pattern mattered for motor abstraction training. The design and procedure were identical to Experiment 7, except that participants completed 5 forward-tracking blocks and 5 backward-tracking blocks instead of rotation-copy blocks. Forward-tracking blocks proceeded as in Experiment 7. In backward-tracking blocks, the sequence of the movement was reversed for the tracking trials. That is, if participants copied the standard motor patterns from point A to point B, they tracked the disc moving from point B to point A during backward-tracking trials.

We preregistered the same sample size and exclusion criteria as for Experiment 7. Two participants were excluded for failing to provide a complete data set and 9 for having too many invalid trials, leaving 89 participants with analyzable data. The comparison between the two types of tracking blocks is shown in Fig. S8.

### Experiments 9–11: Hand-switch generalization

Experiments 9–11 examined whether motor training transfers to canonical performance when training and test movements are executed with different hands. A separate group of participants (*n* = 30 each) completed each experiment using 10 motor patterns, following the design of Experiment 4. Within each block, participants first copied the canonical motor pattern with one hand (pre-training test), then completed three training trials with the opposite hand, and finally copied the canonical motor pattern again with the original hand (post-training test). The three experiments differed only in the content of the three training trials: in Experiment 9, participants copied the canonical (upright) pattern; in Experiment 10, they copied the pattern rotated by three extreme angles (90*^◦^*, 180*^◦^*, 270*^◦^*); and in Experiment 11, they tracked a disc moving along the trajectory of the canonical pattern.

At the start of the experiment, participants reported their dominant hand in a short survey. Hand assignment was counterbalanced within participants across blocks: in half of the blocks, participants tested with the dominant hand and trained with the non-dominant hand, and in the other half this assignment was reversed. To enforce that each phase was performed with the intended hand, participants were required to hold down a set of keys with the non-drawing hand in order to draw: when drawing with the right hand, they pressed Q, W, and E with the left hand, and when drawing with the left hand, they pressed P, O, and I with the right hand. Drawing was disabled unless the correct keys were held, ensuring that the non-drawing hand remained occupied throughout each trial. All other aspects of the procedure, including the demo, copying, and analysis routines, were identical to Experiment 4.

## Data availability

All data and materials are available via OSF at https://osf.io/2gw5c.

## Code availability

All code is available via OSF at https://osf.io/2gw5c.

## Extended Data

**Figure S1:**
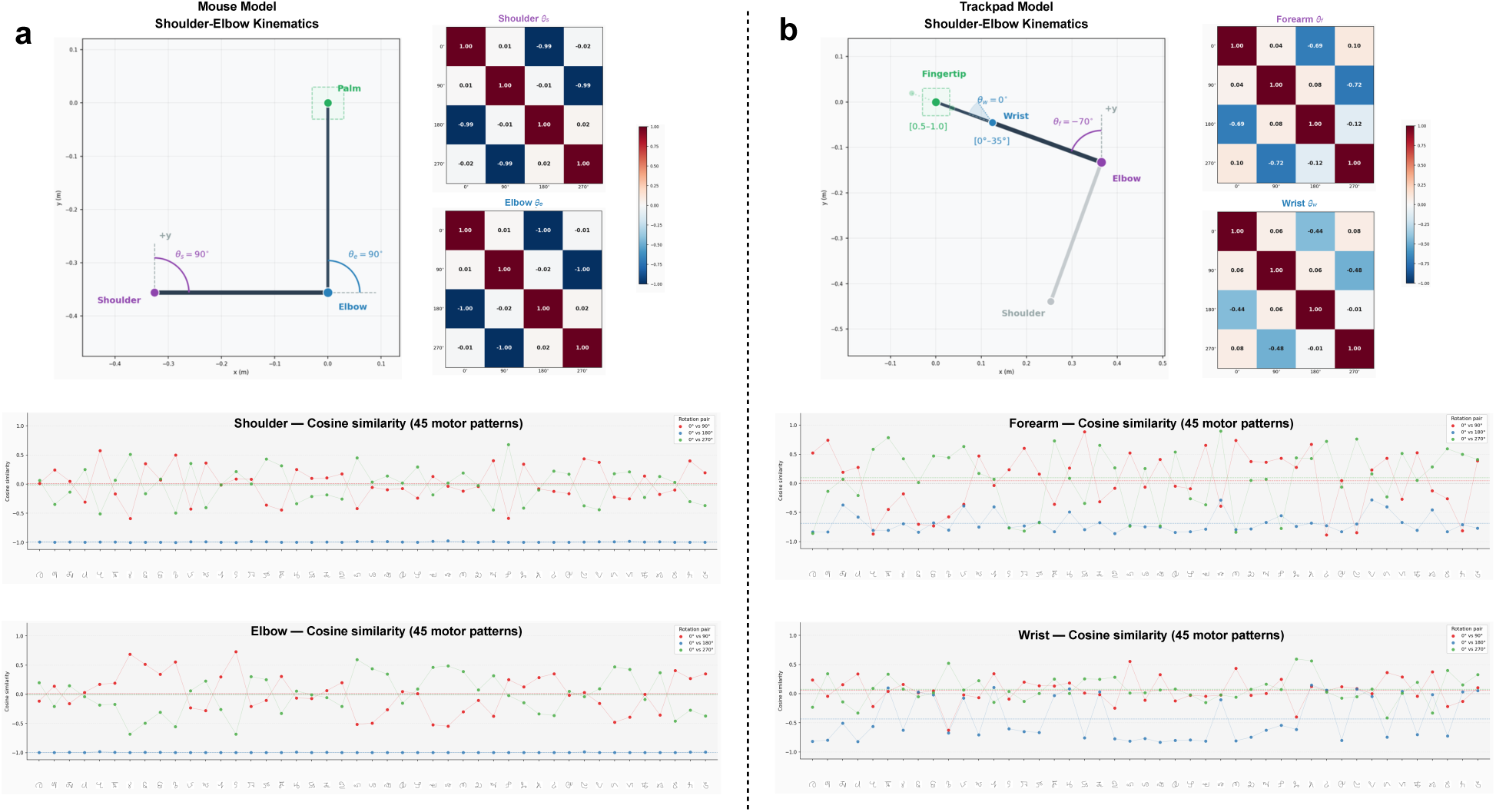
Planar inverse-kinematic models used to estimate joint-angle similarity across rotations. Two effector models were used to test whether rotated motor patterns require similar joint configurations to the canonical (0*^◦^*) patterns. **(a) Mouse model.** A shoulder–elbow–palm chain pivoting at a fixed shoulder, parameterized by the shoulder angle (*θ_s_*) and elbow angle (*θ_e_*). The schematic (top left) shows the initial posture (*θ_s_* = *θ_e_* = 90*^◦^*), with segment lengths drawn to scale. **(b) Trackpad model.** An elbow–wrist–fingertip chain pivoting at a fixed elbow, parameterized by the forearm angle (*θ_f_* ) and wrist angle (*θ_w_*); the initial posture is *θ_f_* = *−*70*^◦^*, *θ_w_* = 0*^◦^*. The gray shoulder–elbow segment is shown for visual context only and is not part of the active kinematic chain. Bracketed values indicate the modeled ranges: wrist angle (0*^◦^*–35*^◦^*) and finger-curl *k*, the ratio of effective to full hand length (0.5–1.0; initial value *k* = 0.70). Within each panel, the two heatmaps (top right) show, separately for each joint angle, the mean cosine similarity between every pair of rotation conditions (0*^◦^*, 90*^◦^*, 180*^◦^*, 270*^◦^*), averaged across all 45 motor patterns; values near 1 indicate near-identical joint trajectories and values near *−*1 indicate opposite (anti-phase) trajectories. The two line graphs (bottom) break these down by pattern, plotting the cosine similarity of each individual pattern between the canonical (0*^◦^*) version and each rotated version (0*^◦^* vs 90*^◦^*, 0*^◦^* vs 180*^◦^*, 0*^◦^* vs 270*^◦^*; see legend). Notably, the cosine similarity between canonical and rotated movement (across 45 characters) does not predict the improvement of performance (mouse model: r = 0.16, p = 0.29; trackpad model: r = -0.067, p = 0.66), suggesting that the improvement is not driven by the low-level movement similarity between canonical and rotated patterns. See the full detail of the model and visualization at https://zk.actlabresearch.org/abstraction/movement_models.html

**Figure S2:**
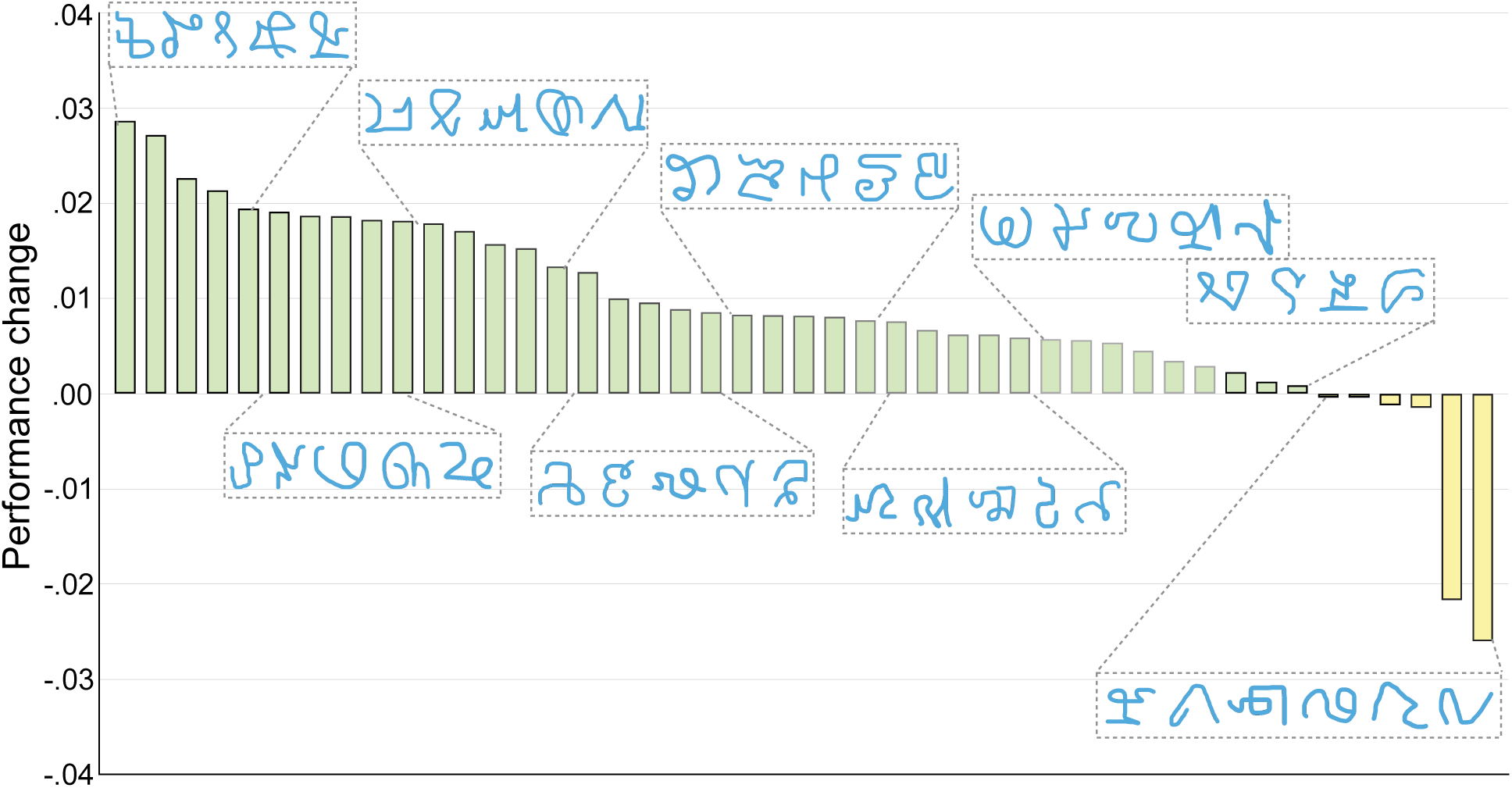
Abstraction training produced performance improvement across the large majority of individual motor patterns. The performance index change (Trial 5 minus Trial 1) is shown separately for each of the 45 motor patterns learned by participants in Experiment 1. Positive values indicate improvement following abstraction training. The majority of patterns (39 out of 45; 87%) showed a positive training effect, demonstrating that the learning benefit of abstraction training generalizes broadly across motor patterns rather than being driven by a small subset of easily learned items.

**Figure S3:**
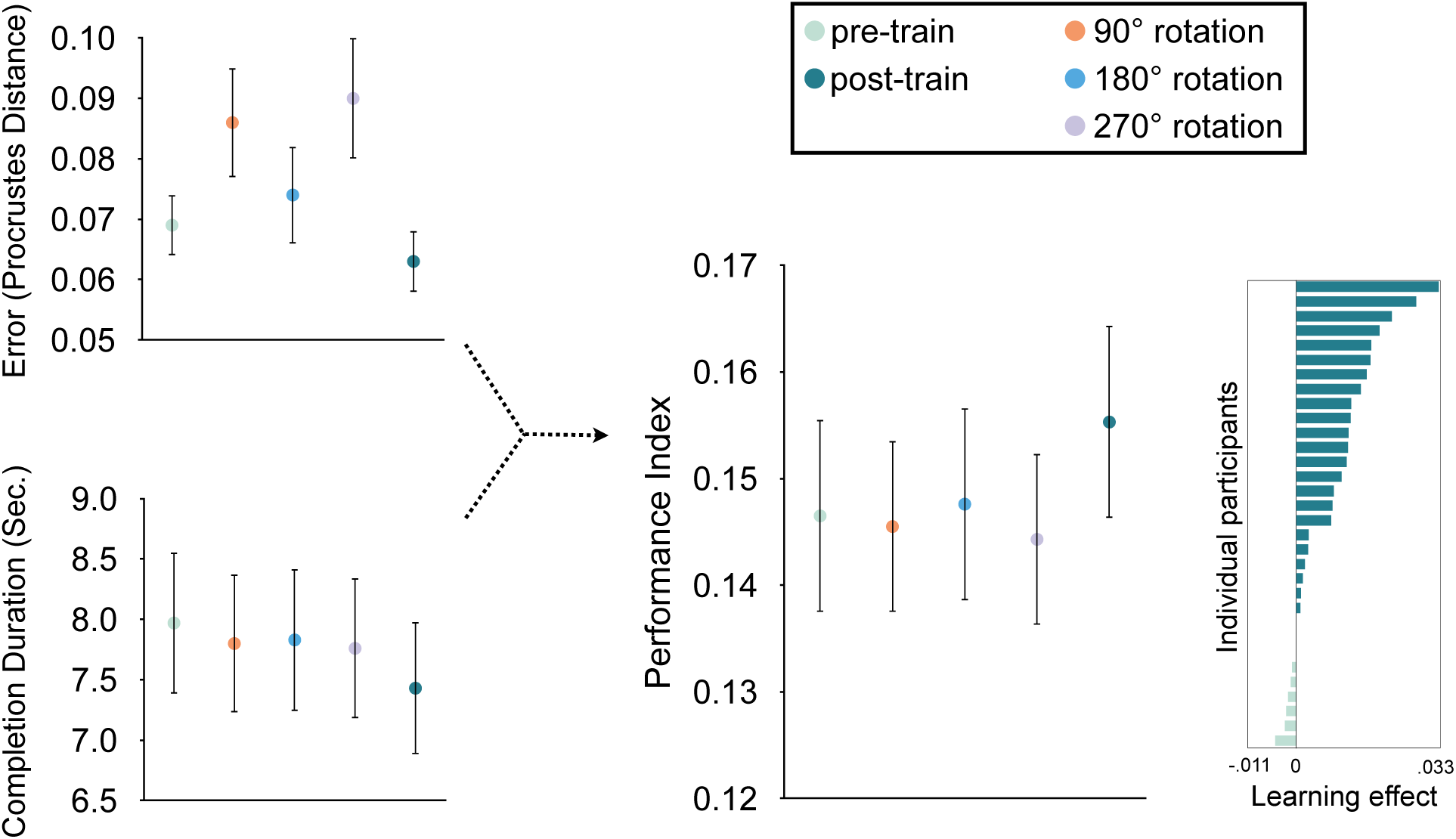
Performance changed across the learning block and was consistent across participants in Experiment 1. (**a**) Mean trajectory error (Procrustes distance), movement time, and performance index across all five trials of the learning block, averaged across participants and motor patterns in Experiment 1. Error bars represent s.e.m. (**b**) Individual performance change scores (Trial 5 minus Trial 1 performance index) for all 30 participants. Positive values indicate improvement following abstraction training. The large majority of participants showed performance improvement, confirming that the group-level result reflects consistent improvement across individuals rather than an outlier-driven effect.

**Figure S4:**
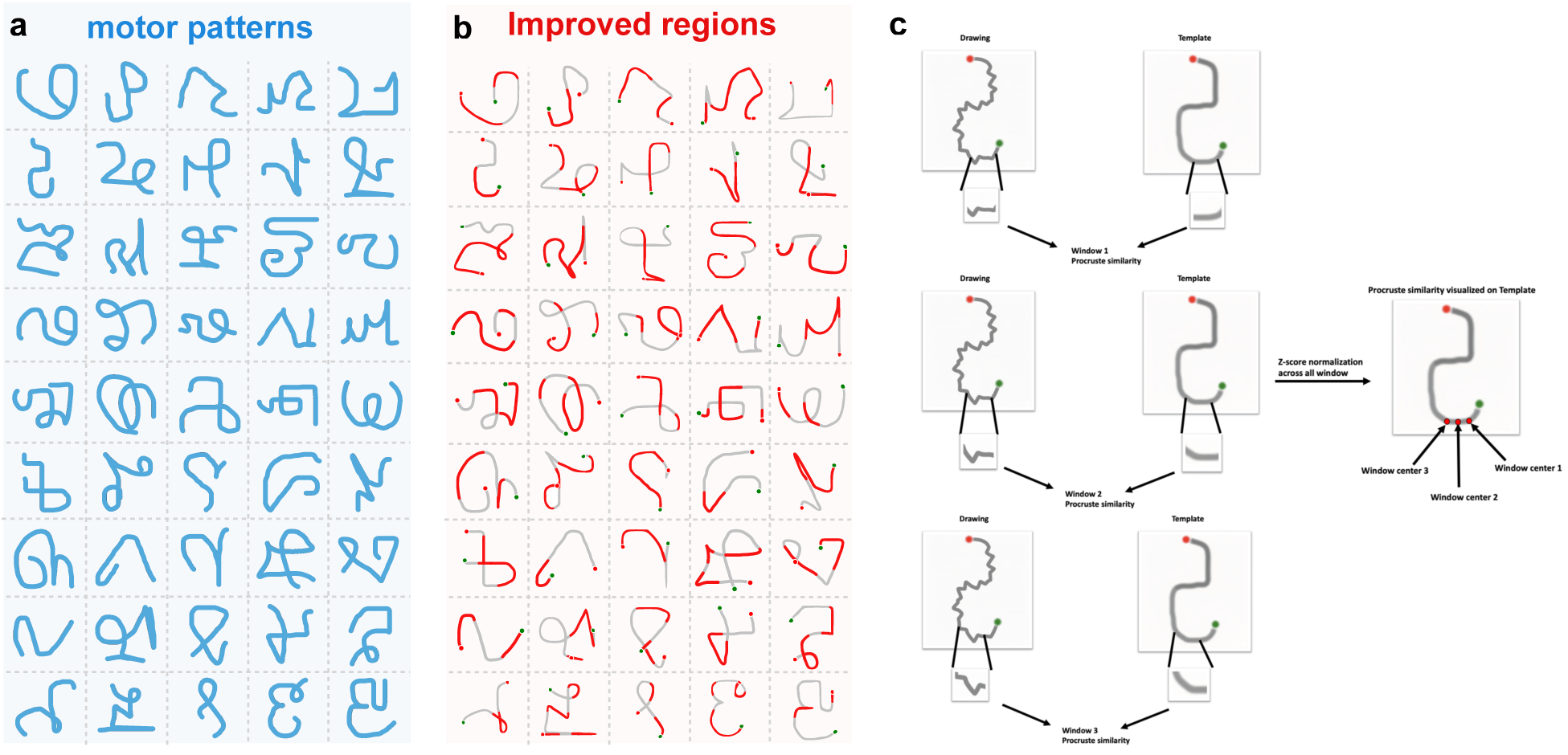
Abstraction training selectively improves accuracy at specific segments of each motor pattern. (**a**) The set of motor patterns used in Experiment 1. (**b**) Improved segments for each motor pattern following abstraction training, identified using a sliding window analysis. Warmer colors indicate segments showing greater accuracy improvement from Trial 1 to Trial 5. (**c**) Schematic illustration of the sliding window analysis procedure. A window of 30 trajectory points was slid along each motor pattern trajectory with a stride of 29 points. Both the template and drawn trajectories were resampled to a common length prior to analysis, enabling direct Procrustes distance comparison within each window. For each motor pattern, Procrustes similarity values across all windows were *Z*-score normalized, and the resulting *Z*-scores were mapped onto the spatial coordinates of the template trajectory to identify regions of greatest improvement.

**Figure S5:**
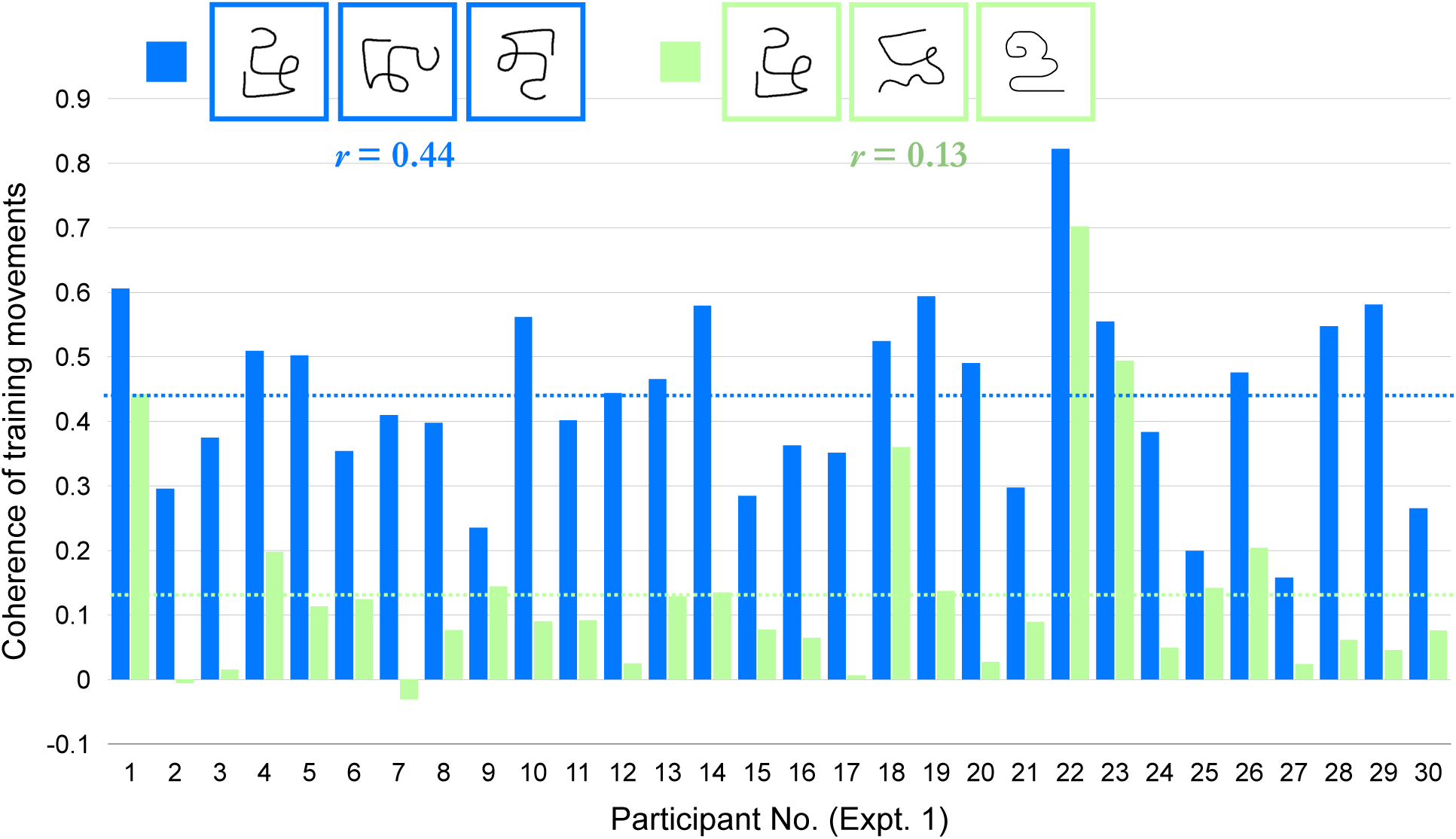
Rotation training preserves pattern-specific temporal dynamics, though temporal coherence does not predict learning magnitude. Each motor pattern was segmented into 10 equal-distance constituents, and the proportion of time spent traversing each segment was computed for each training trial. Temporal coherence was defined as the correlation between the temporal dynamics — the relative time spent on each segment — across the three rotation training trials within a block. Temporal coherence during rotation training was compared to temporal coherence computed across trials in which participants drew different, unrelated motor patterns. Participants showed significantly higher temporal coherence during rotation training than across different-pattern trials, indicating that consistent, pattern-specific temporal dynamics were preserved even when the spatial structure of the movement was dramatically altered by rotation. However, individual differences in temporal coherence did not predict the magnitude of the training effect, suggesting that while pattern-specific temporal structure is present during abstraction training, it is not the primary driver of learning.

**Figure S6:**
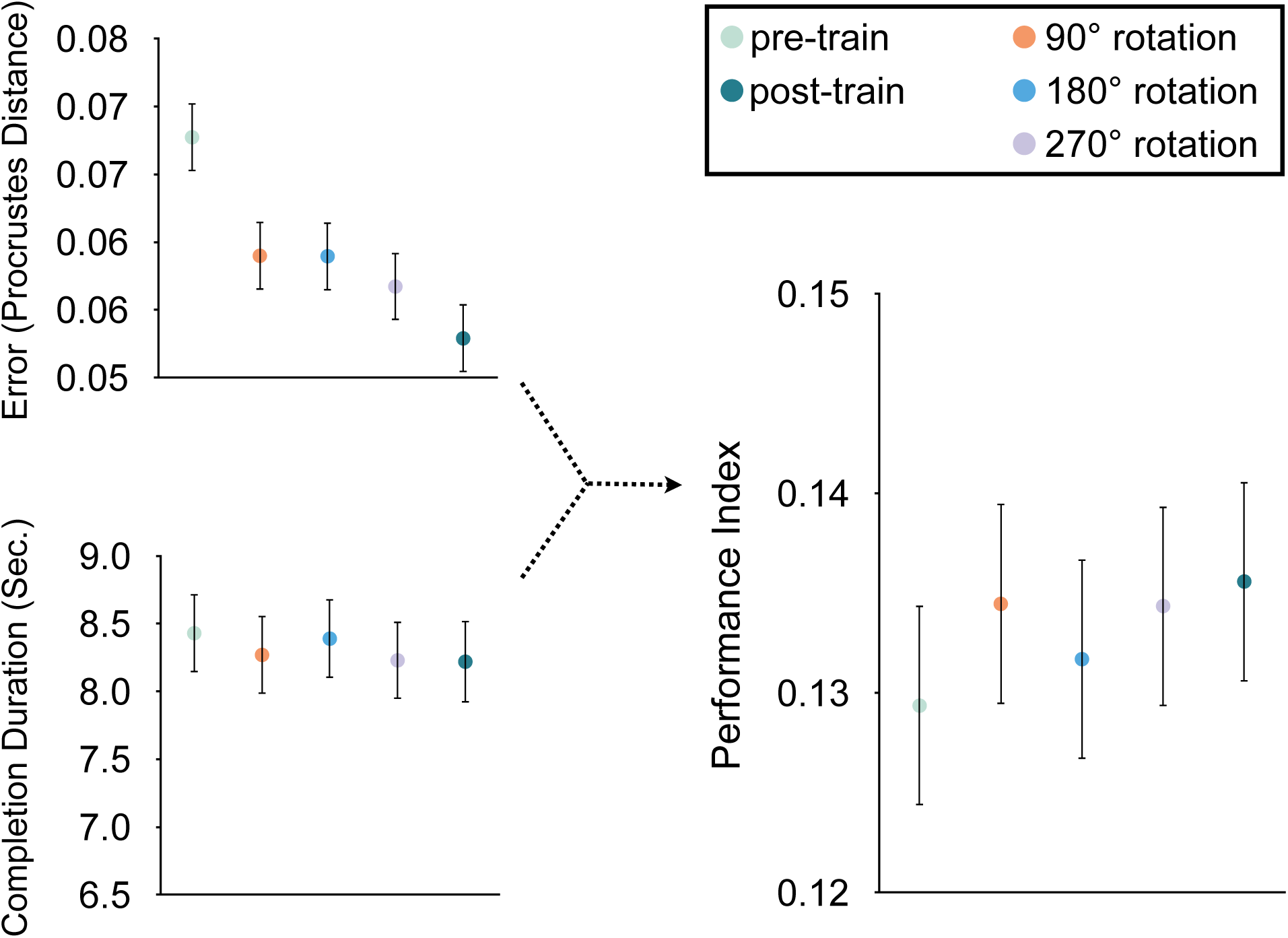
Aphantasic participants showed progressive performance improvement across the learning block in Experiment 4. Mean trajectory error (Procrustes distance), movement time, and performance index across all five trials of the learning block, averaged across participants and motor patterns in Experiment 4 (aphantasic sample; *n* = 88). Error bars represent s.e.m. The pattern of improvement across trials in aphantasic participants closely mirrors that observed in neurotypical participants in Experiment 1, providing further evidence that the learning benefit of abstraction training does not depend on the capacity for conscious visual mental imagery.

**Figure S7:**
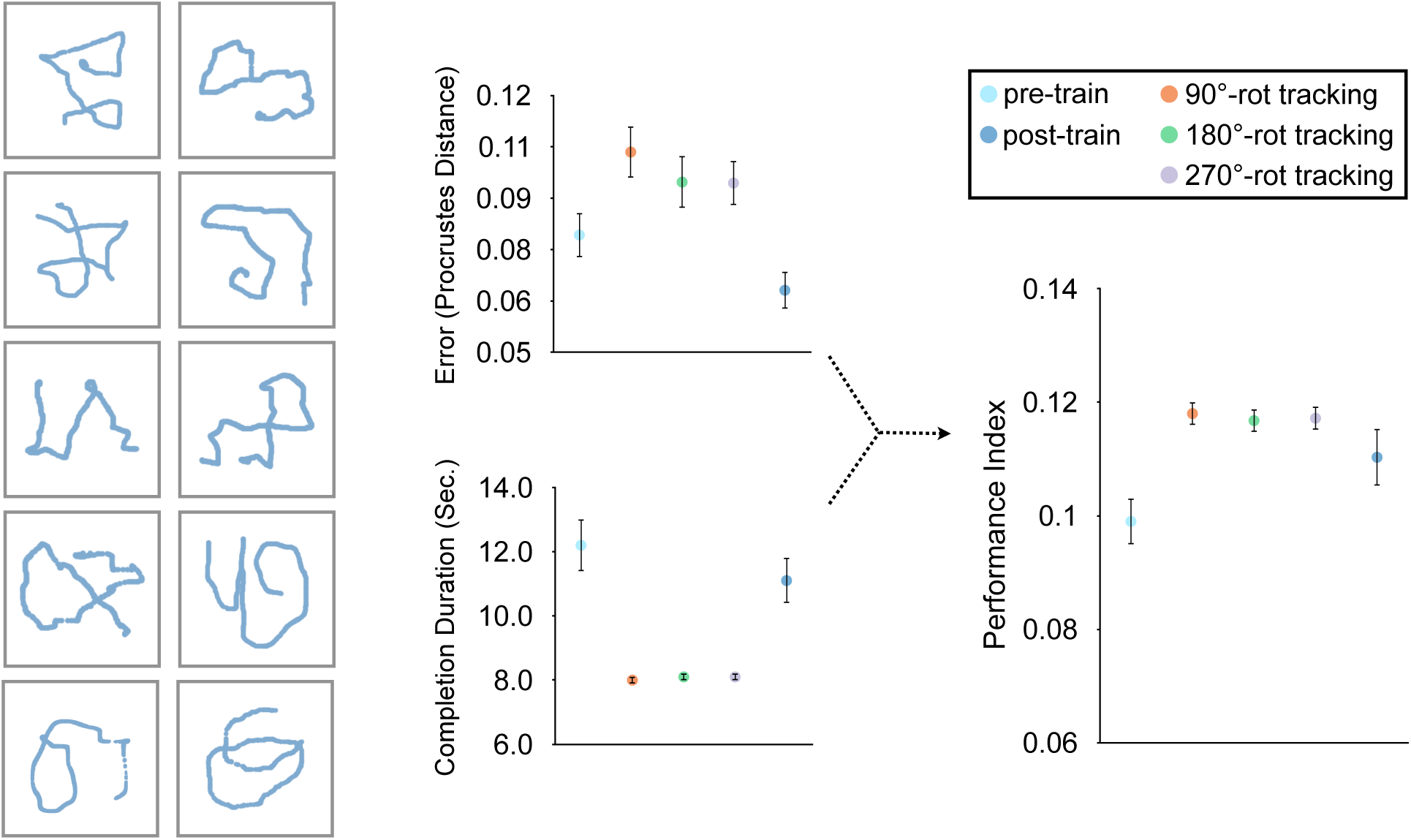
Tracking training produced systematic improvement across the learning block in Experiment 7. (**left**) Sample movement trajectories produced by participants during tracking training trials, illustrating the quality of tracking performance across the rotated motor pattern trajectories. (**right**) Mean trajectory error (Procrustes distance), movement time, and performance index across all five trials of the learning block for both abstraction training blocks and tracking training blocks, averaged across participants and motor patterns in Experiment 7 (*n* = 87). Error bars represent s.e.m. Performance on the canonical motor pattern improved progressively across the block in both training conditions, with tracking training producing a stronger improvement than standard abstraction training by the final trial.

**Figure S8:**
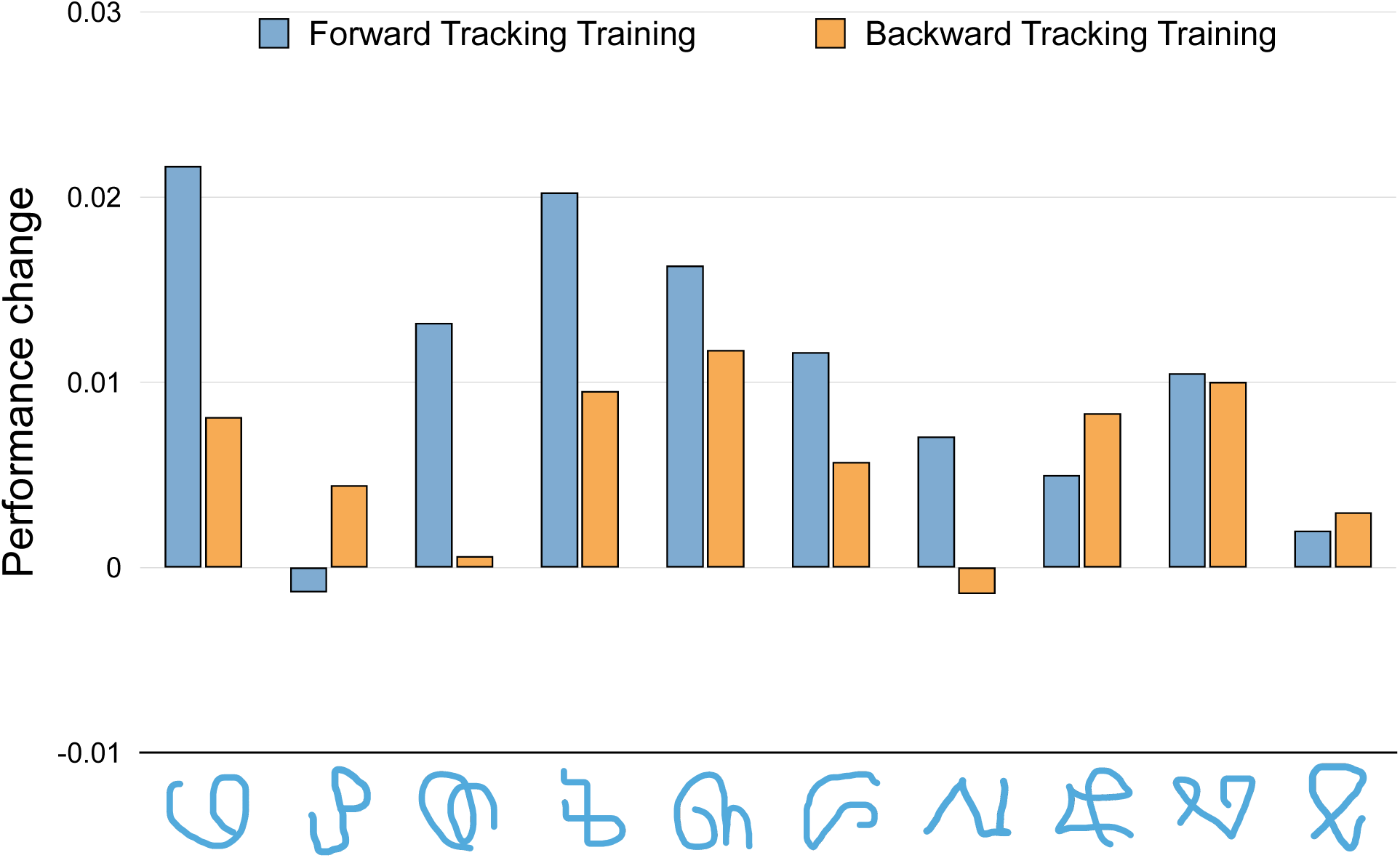
The advantage of forward over backward tracking was varied across individual motor patterns in Experiment 8. Performance index change (Trial 5 minus Trial 1) following forward tracking training and backward tracking training is shown separately for each of the 10 motor patterns used in Experiment 8. Forward tracking produced a larger learning effect than backward tracking for the majority of individual motor patterns, demonstrating that the sequential order advantage observed at the group level reflects a consistent pattern across items rather than being driven by a subset of patterns with atypical trajectory structure.

**Figure S9:**
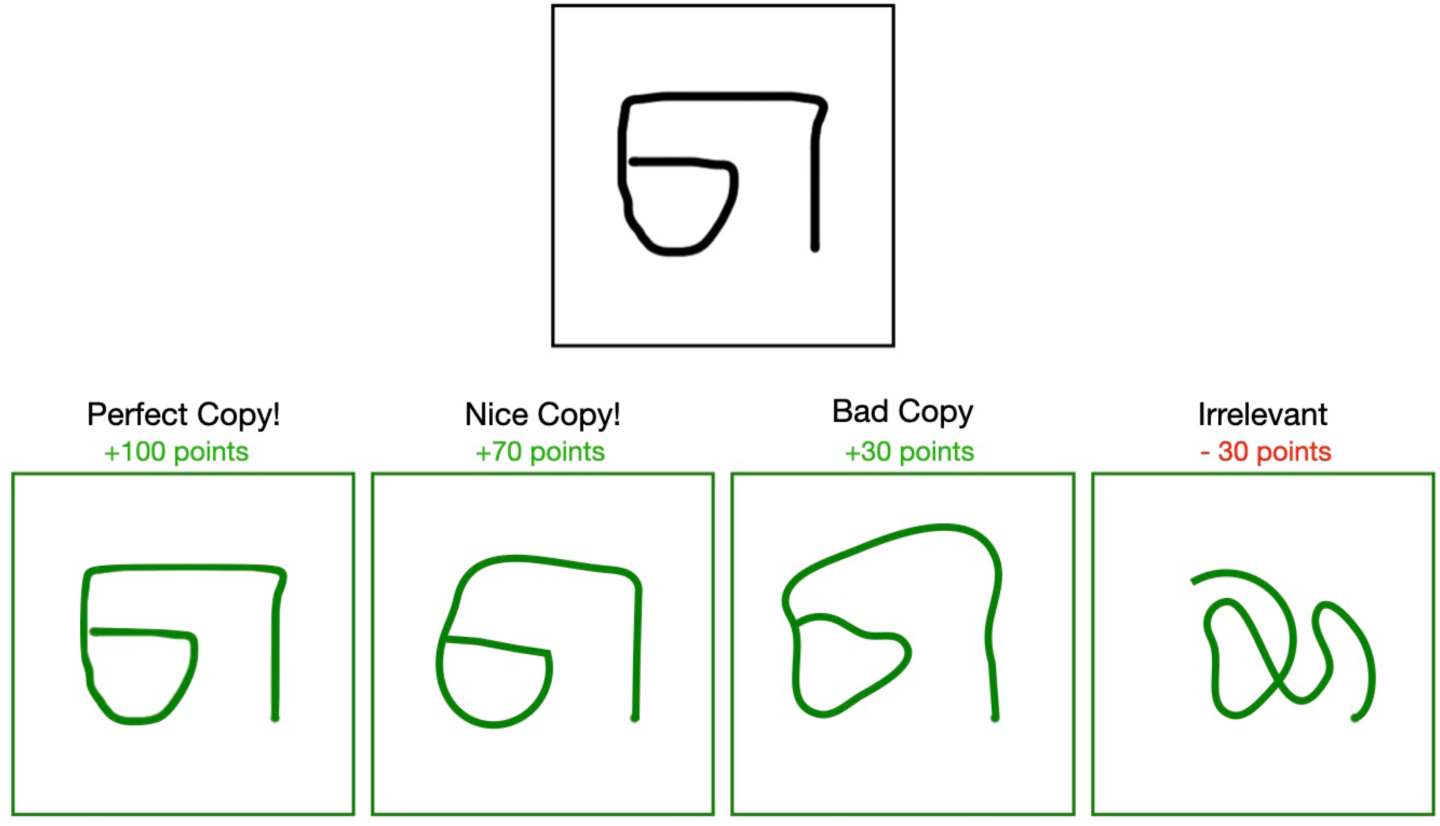
Score sheet provided to participants as part of the experimental instructions.

